# Optogenetic and sonogenetic pacing of unmodified cardiac tissue upon quick integration with specialized biological actuators

**DOI:** 10.64898/2026.10.19.760310

**Authors:** Morgan E. Pettebone, Julie L. Han, Amin Forouzandehmehr, Olivia Chesla, Christianne J. Chua, Maria R. Pozo, Yuli W. Heinson, Wei Liu, Emilia Entcheva

## Abstract

Electrical stimulation for cardiac pacing has been in clinical use for 100 years and it is the current standard. Alternative scalable pacing methods that are contactless, with good spatial precision and can also be cell-selective are of interest for both *in vitro* and *in vivo* applications. Cardiac pacing by light or by ultrasound (US) can be achieved using optogenetic transduction of the tissue and by enhancing responsiveness to ultrasound. In this work we create specialized biological actuators (cell spheroids) out of generic human cells, modified optogenetically (using Channelrhodopsin2) and sonogenetically (using human Piezo1), guide them to a desired location to promote quick integration with unmodified cardiac syncytia within less than two hours. We quantify their ability to confer pacing by light or US and to entrain the native cardiac tissue, modeled here with human induced pluripotent stem cell derived cardiomyocytes (hiPSC-CMs) grown in monolayers, quasi-3D oriented microtissues or cell spheroids. Magnetic guidance (via magnetic nanoparticles) speeds up integration and facilitates positioning to optimize electrotonic load for pacing; size match and three-dimensional growth also accelerate integration. Computational modeling corroborates the possibility to pace by light and by US. Using optogenetically-gated US pulses (0.5 MHz), we show entrainment in voltage and calcium responses and identify the best performing configurations. Experiments reveal the potential role of Piezo1, esp. when activated with Yoda1. The approach is shown to work with physiologically relevant heart cells, such as human cardiac fibroblasts as optimized actuators, and therefore can be considered for *in vivo* applications as part of heart regeneration efforts. High-throughput cardiotoxicity testing and drug screening *in vitro* can directly benefit from deployment of such biological actuators.

## INTRODUCTION

Cardiac pacemakers epitomize a successful and impactful technology that continues to evolve and improve. Nevertheless, there are scenarios where alternative solutions to the clinically used electrical pacemakers are of interest. Such scenarios include temporary “bridge” pacing[1], potential temporary entrainment and phase resetting during regeneration efforts[2, 3], pre-procedure external probing of activation patterns for arrhythmia control [4, 5] or the search for more physiologically-adaptable solutions [6]. Various experimental approaches for cardiac pacemaking have been developed over the last two decades. Biological pacemakers have been pursued using gene and cell engineering, including transcriptional reprogramming based on TBX18 [7] or various approaches involving the genes underlying pacemaking currents, e.g. HCN2 [8–10], as well as the design of a fully bioengineered sinoatrial node[11]. Progress in materials science and miniaturization yielded bioresorbable versions of minimally invasive pacemakers[1]. Optogenetics opened the possibility to explore light-based technologies for pacemaking [12, 13] after genetically expressing light-sensitive ion channels (opsins) in the heart [14–19]. Due to limitation of light penetration depth, alternative modalities have been explored, including direct ultrasound (US) based pacing [5, 20–23] and sonogenetic pacing (after genetic sensitization of cardiac tissue to mechanical stimulation)[24, 25]. Some of these approaches have limitations, including gene therapy challenges (faced by biological pacemakers using transcriptional reprogramming, optogenetic and sonogenetic pacemakers), specificity of targeting (ultrasound pacing), non-biological resorption by-products (man-made bioresorbable pacemakers).

In this report we explore the fundamental possibility to design small scalable biological structures endowed with specific responsiveness to external energies (light, ultrasound) that can quickly couple to native cardiac tissue and be remotely controllable to achieve pacemaking. These can be built from patient-derived cells to minimize immune responses, can be introduced using cell delivery methods and, after integration, can be engaged on demand with the specific energy modality. Previously, we introduced the tandem-cell-unit (TCU) concept to optogenetic pacing, where a non-excitable cell or structure, sensitized to light, can couple to unmodified cardiac syncytia and be used to control its rhythm[3, 19, 26]. In this approach, the bioactuator (sensitized to light or other energy modality) translates a pulse of that energy into a sharp change in membrane voltage through the induced ion current(s), which in turn triggers the electric event in the unmodified target cells/tissues, if they are coupled to the bioactuator[19]. Here we demonstrate the broader applicability of the TCU concept not only to optogenetic but also to sonogenetic biological drivers for pacemaking. This study focuses on early integration, scalability, precise positioning and enhanced contact (using magnetic guidance), and generalizability to different cell types. This proof-of-principle *in vitro* investigation, supported by computational modeling, quantifies various key factors determining quick integration and entrainment of the native tissue, modeled here by syncytia of human induced pluripotent stem cell derived cardiomyocytes (hiPSC-CMs). As part of this project, we developed a novel integrated platform that allows for optogenetic and ultrasound pacing while optically monitoring voltage and calcium in the various studied 2D and 3D hiPSC-CM constructs, as well as a modified in-incubator optical actuation and sensing system to track entrainment. Computational modeling of optogenetic and sonogenetic stimulation of hiPSC-CMs accompanies our experimental results. The approach can be immediately impactful *in vitro* for remote scalable actuation in high-throughput platforms for drug testing and can be potentially integrated in cell delivery approaches, currently in early clinical trials for cardiac regeneration[27–29].

## RESULTS

Building upon prior work using the TCU approach for optically-controllable cardiac syncytia [19, 26], we aimed to leverage multiple modes of actuation, namely optical and mechanical, and to quickly entrain both 2D and 3D cardiac structures. Precise positioning and contact enforcement was done using magnetic nanoparticles (mNPs). The main objective was to uncover the factors contributing to rapid and robust cardiac entrainment, with potential applications in areas like HT-drug screening, tissue engineering and regeneration (**Fig. 1**).

**Figure 1.**
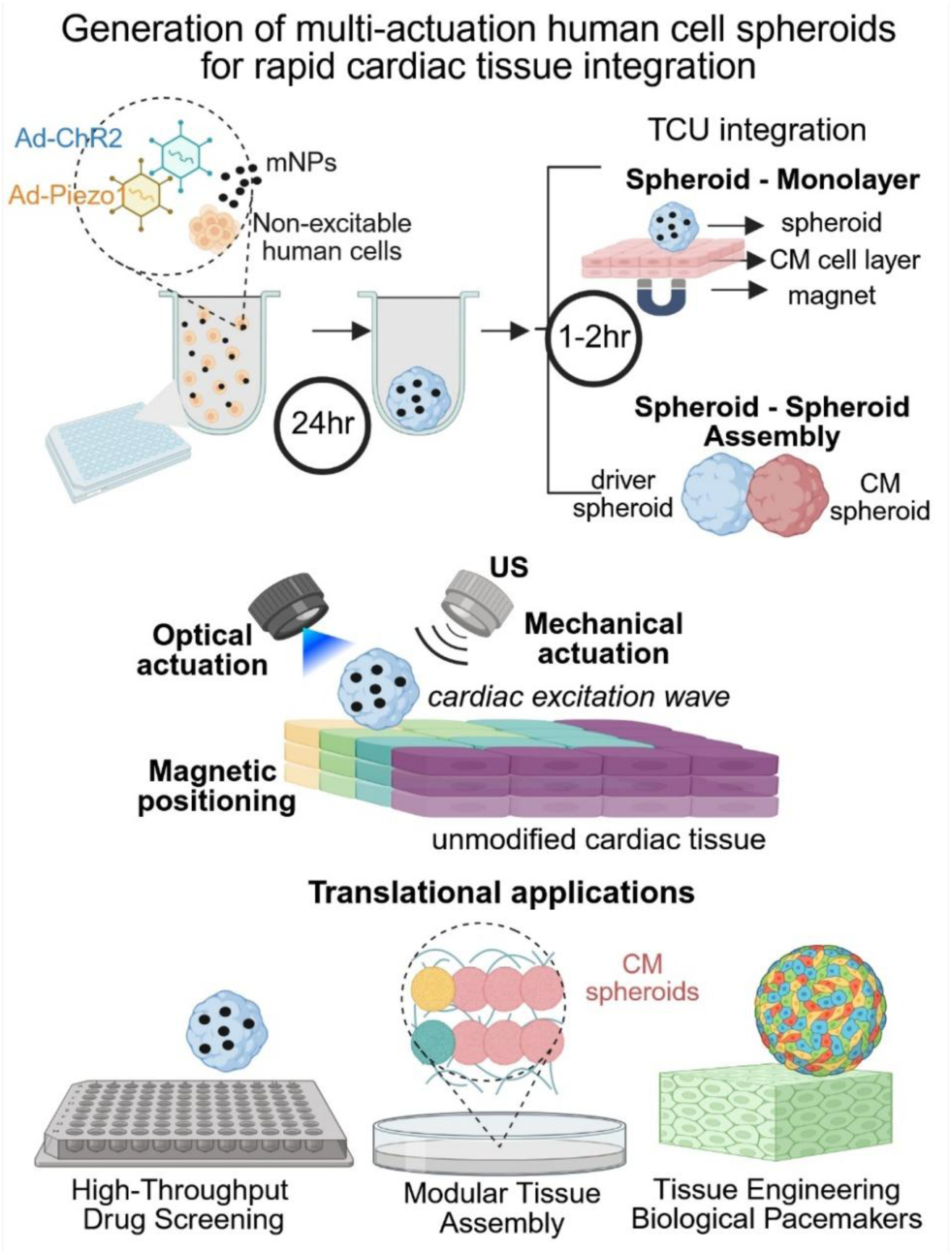
Overview of the generation, integration and deployment of cell spheroids for actuation of unmodified cardiac tissue. Cell spheroids can be generated using generic non-excitable human cells, genetically sensitized to respond to light (optogenetic modification with ChR2) or ultrasound (via Piezo1), with the addition of magnetic nanoparticles, mNPs, to allow magnetic guidance and positioning. Quick integration (1-2 hours) of the assembled spheroids can be achieved with unmodified cardiomyocyte syncytia (helped by magnetic positiong) or with unmodified human cardiomyocyte spheroids or organoids. As per the tandem-cell-unit (TCU) approach, the resultant assemblies can be easily stimulated electrically after quick integration using light or ultrasound. Potential future applications include HT studies, modular tissue assembly, engineering of biological pacemakers and magnetic navigation and actuation *in vivo*.

### Rapid and robust control by optogenetic stimulation of unmodified cardiac tissues using magnetic nanoparticles and 3D scaffolding

Previous work focused on characterizing Channelrhodopsin2 (ChR2)-expressing human embryonic kidney cells (ChR2-HEK) cell spheroids as a convenient model system to demonstrate their potential use as actuators on cardiac cell monolayers[26]. Here we focused on optimizing early integration (within two hours from contact). ChR2-HEK cells were loaded with mNPs overnight before growing cell spheroids and depositing these onto unmodified hiPSC-CMs monolayers in high-throughput (HT) 96 well plates. A magnetic plate underneath was used to guide and aid the quick positioning of the ChR2-HEK spheroids, **Fig. 2A**. Entrainment (1:1 responses to pacing) within the first 1-2 hours was graphed for samples with spheroids +mNPs and those without mNPs, **Fig. 2B**. We defined a sample as entrained if it was able to follow light pulses at a 1:1 beat ratio for at least 8 seconds for at least 2 pacing frequencies. In this early time interval, only a fraction of the samples without mNPs (7/16, or 44%) paced, while the entrainment rate improved for +mNPs spheroids (15/22, or 68%), **Fig. 2C**. Furthermore, the samples with spheroids +mNPs guided to the edge of a monolayer showed increased entrainment, with 70% (17/24) being entrained compared to just 25% (4/16) being entrained if the spheroid was deliberately positioned away from the monolayer edge, **Fig. 2C**. This suggests that spheroid positioning onto a flat syncytium is a main attribute for drivability regardless of mNP presence; magnetic guidance can greatly aid in positioning of the spheroid for effective pacing. An edge position reduces the “electronic load” or the immediate area that the spheroid needs to excite. Multiple sizes and types of mNPs were tried to see the effect on spheroid controllability and cardiac entrainment, without significant differences in entrainment ranges between the tested mNP types, **Fig. S1**.

**Figure 2.**
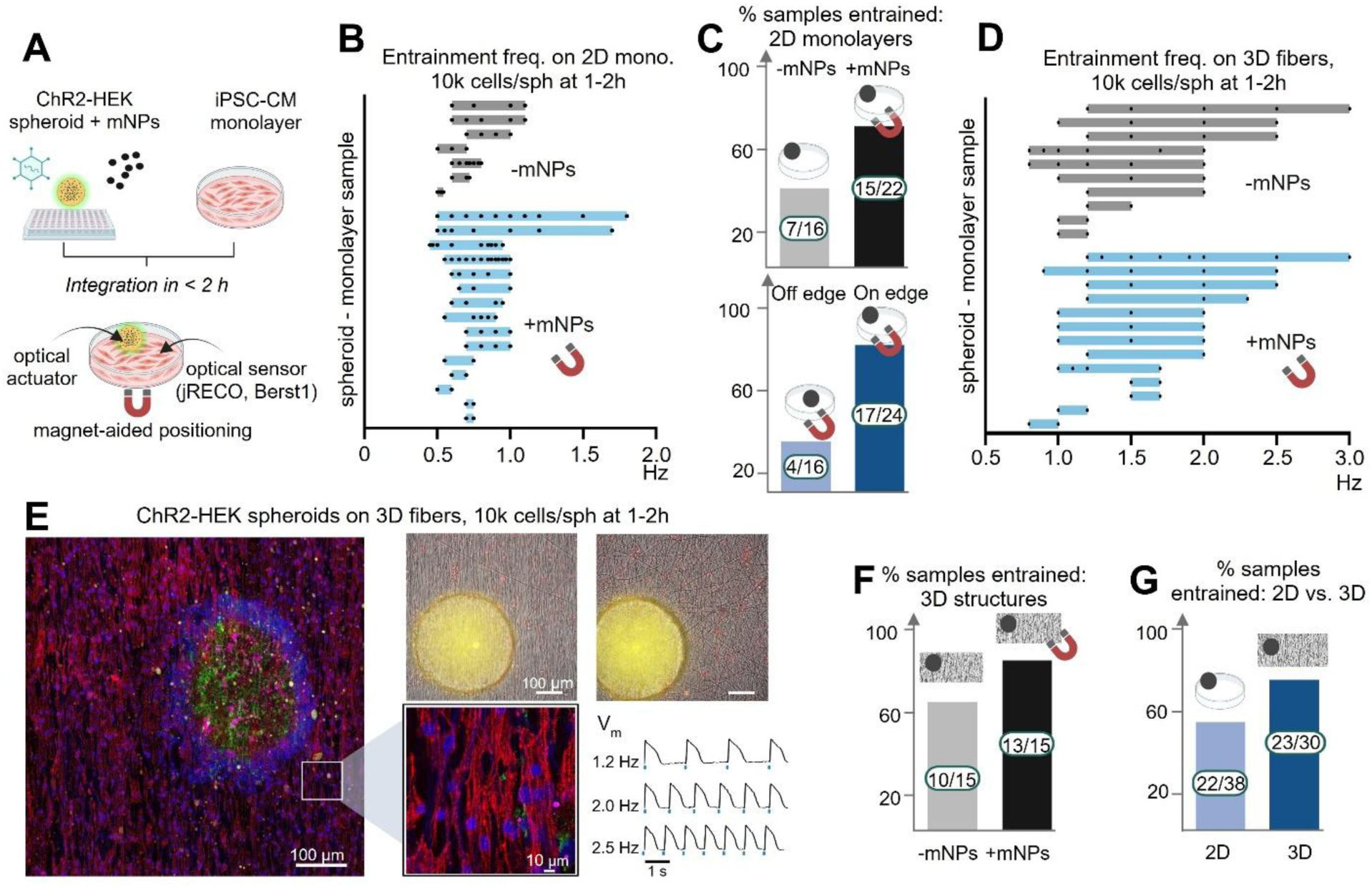
Magnetic guidance and 3D scaffolding improve rapid optogenetic entrainment of unmodified cardiac tissue. **A.** Experimental setup for probing entrainment of human iPSC-CM monolayers by ChR2-HEK spheroids +/- mNPs. The addition of mNPs to spheroids can aid in actuator placement and integration. **B.** Entrainment ranges from 2D iPSC-CM monolayers with (n=16) and without (n=22) mNPs (Takara) added to the spheroids (10k cells) after 1-2h integration. Only samples following at least two pacing frequencies are shown. **C.** Top: Percentage of 2D monolayer samples entrained -/+ mNPs and magnetic pull. Bottom: Percentage of 2D monolayers entrained after magnetic positioning of the spheroid (on edge vs. center). Takara and Micromod mNPs were used. **D.** Entrainment ranges from iPSC-CMs grown on 3D fibers with (n=15) and without (n=15) mNPs (Micromod) added to the spheroids (10k cells) after 1-2h integration. Only samples following at least two pacing frequencies are shown. **E.** Left: Confocal image of ChR2-HEK spheroid placed on an iPSC-CM syncytium grown on electrospun nanofiber scaffold. Red shows alpha-actinin, blue shows nuclei, green shows ChR2-eYFP, and pink shows mNPs (Micromod) tagged with mCherry. An inset of the confocal image shows the aligned cardiomyocytes with alpha-actinin structure in greater detail. Right, top: Images showing ChR2-HEK spheroids placed on locally aligned iPSC-CM samples in brightfield overlayed with red (voltage-sensitive BeRST1 dye) and yellow (eYFP-ChR2). Right, bottom: Example voltage entrainment data collected from an aligned iPSC-CM sample. **F.** Percentage of 3D cardiac structures entrained by spheroids -/+ mNPs. **G.** Comparison of 2D vs. 3D cardiomyocyte syncytia entrained by spheroids, paced optogenetically.

To test how entrainment would be impacted by more realistic 3D structural cues from cardiac tissue, hiPSC-CMs were plated on locally aligned electrospun nanofibers, **Fig. 2E**. Their entrainment ranges were subsequently probed after adding ChR2-HEK spheroids both with and without mNPs, **Fig. 2D**. This 3D growth improved entrainment for both groups - (-mNPs: 66% vs. +mNPs: 87%), **Fig. 2F**. Samples showed greatly increased frequency pacing ranges on 3D vs. 2D structures (**Figs. 2B, 2D**), with overall increase in drivable samples in 3D to 76% (23/30) vs. 58% (22/38) for 2D monolayers, regardless of magnetic guidance, **Fig. 2G**. This improved integration and entrainment is likely due to the naturally increased contact area and “lodging” of the pacing spheroids within the 3D topography – resembling interaction with native tissue.

### Optimized optogenetic entrainment of spheroid-spheroid assemblies

To further explore the role of 3D interactions that increase contact area and as a step towards “assembloid” formation[11, 30], we studied hybrid spheroid-spheroid pairs/assemblies. ChR2-HEK spheroids were manufactured, without mNPs. In addition to growth in standard (96 and 384 well) spheroid plates, we tested mass production using EZ-Sphere to produce up to 2,700 optogenetic spheroids with a single cell deposition and characterized their resulting sizes after 1 day, providing a method for scaling up production. Human iPSC-CM spheroids were produced and grown for 7-9 days before pairing. We prelabeled the CM spheroids with a voltage-sensitive fluorescent dye and examined the immediate pairing of the two spheroids, **Fig. 3A**. Our study probed how different aspects, e.g. spheroid size, size matching and integration time, impacted the resulting entrainment ranges. Insufficient integration time (less than one hour) produced inconsistent entrainment ranges (**Fig. S2**) while larger spheroid sizes caused a reduction in overall entrainment ranges (**Fig. S3**). Cell spheroid pairs of 1000 cells each, when integrated over at least 2 hours, provided the most consistent entrainment and the responses with the widest frequency ranges (**Fig. 3B**) while being large enough to see and handle relatively easily. We found that size matching the spheroid-spheroid pairs also greatly impacted entrainment, **Fig. 3B**. Overall, the spheroid-spheroid entrainment was much easier than spheroid-monolayer integration (likely due to the natural ability to merge and interact in 3D), yet the delicate spheroid pairing process – ensuring contact – was important.

**Figure 3.**
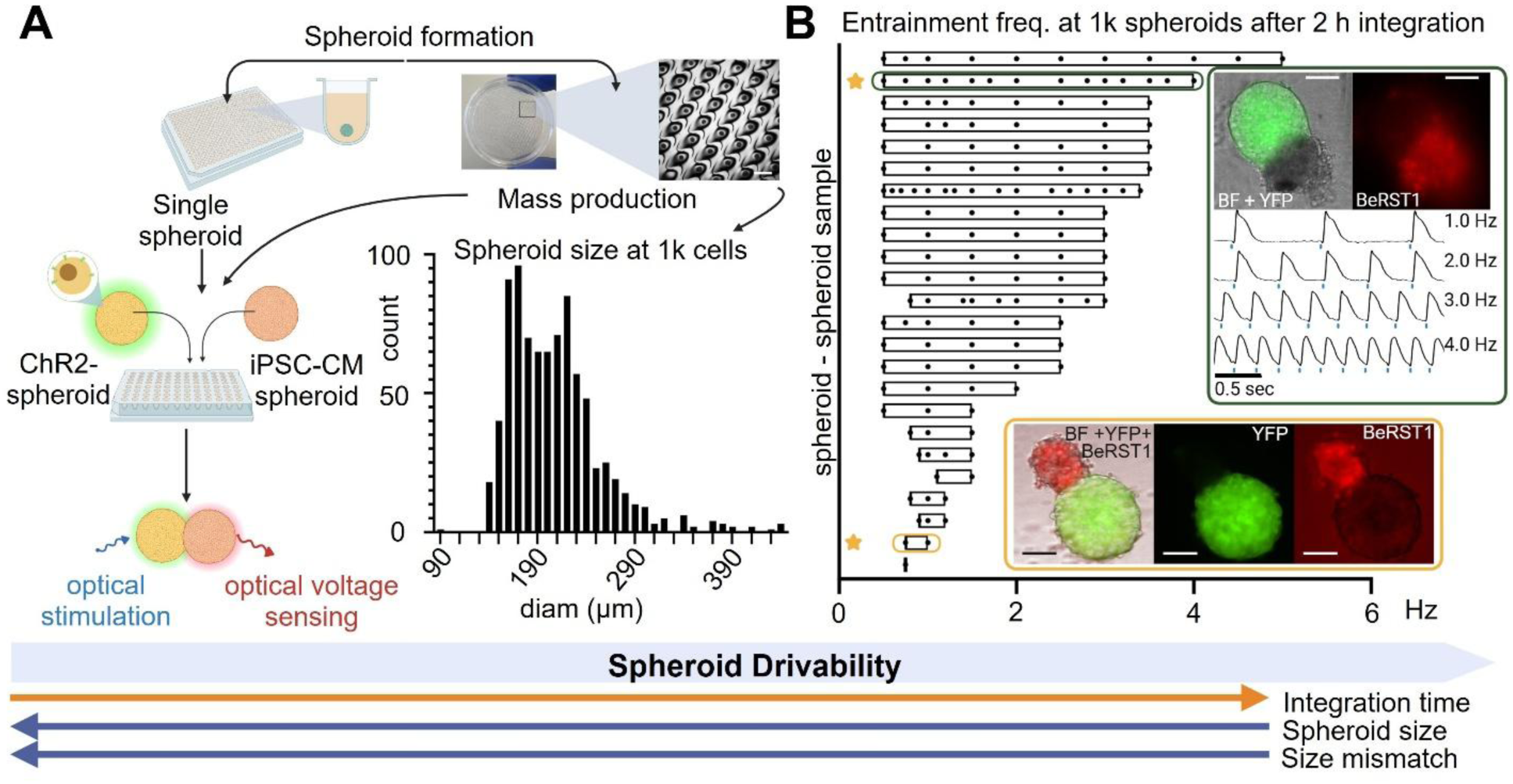
Optogenetic entrainment of spheroid-spheroid assemblies: ChR2-HEK spheroids can robustly drive unmodified cardiac spheroids when well-matched in size and after adequate integration time. **A.** Spheroid generation methods using either spheroid plates, for individual spheroid formation per well, or an EZ-Sphere dish for mass production. A histogram of spheroid sizes is included for characterization of ChR2-HEK spheroids seeded at 1,000 cells/well in the 35-mm EZ-Sphere dish. The ChR2-spheroid is then combined with a iPSC-CM spheroid in a single well for 1-2 hours until optical actuation is achievable. **B.** Entrainment frequency ranges for 1k cell ChR2-HEK and unmodified iPSC-CM spheroid pairs after 2 hours integration. The top inset shows example images and voltage traces for a robustly-pacing and well size-matched pair, while the bottom inset highlights a pair with poorer entrainment range due to significant size mismatch. Samples that did not pace not shown. Scale bars are 100 µm.

### Optogenetic human cardiac fibroblast (cFB) spheroids and their automated characterization

We sought to generalize the TCU approach to optogenetic actuation with spheroids formed by cardiac-relevant cell type. Cardiac fibroblasts, cFBs, being one of the most abundant cell types in the human heart, were a natural first option. In previous studies we have demonstrated optogenetic pacing through co-cultures of ChR2-expressing primary cardiac fibroblasts and cardiac myocytes[31–34]. To check if ChR2 expression affects the spheroid formation and growth of our adult human cFBs, here we used an Etaluma LS850 microscope for automated tracking over multiple days, inside a standard incubator. We found similar size and growth patterns over time between ChR2-cFBs and non-infected human cFBs, for matching seeding cell density, **Figs. 4**. Robust ChR2 expression was achieved after optimization. These results confirmed that ChR2-cFBs are good candidates for optogenetic biological actuators using the TCU approach.

**Figure 4.**
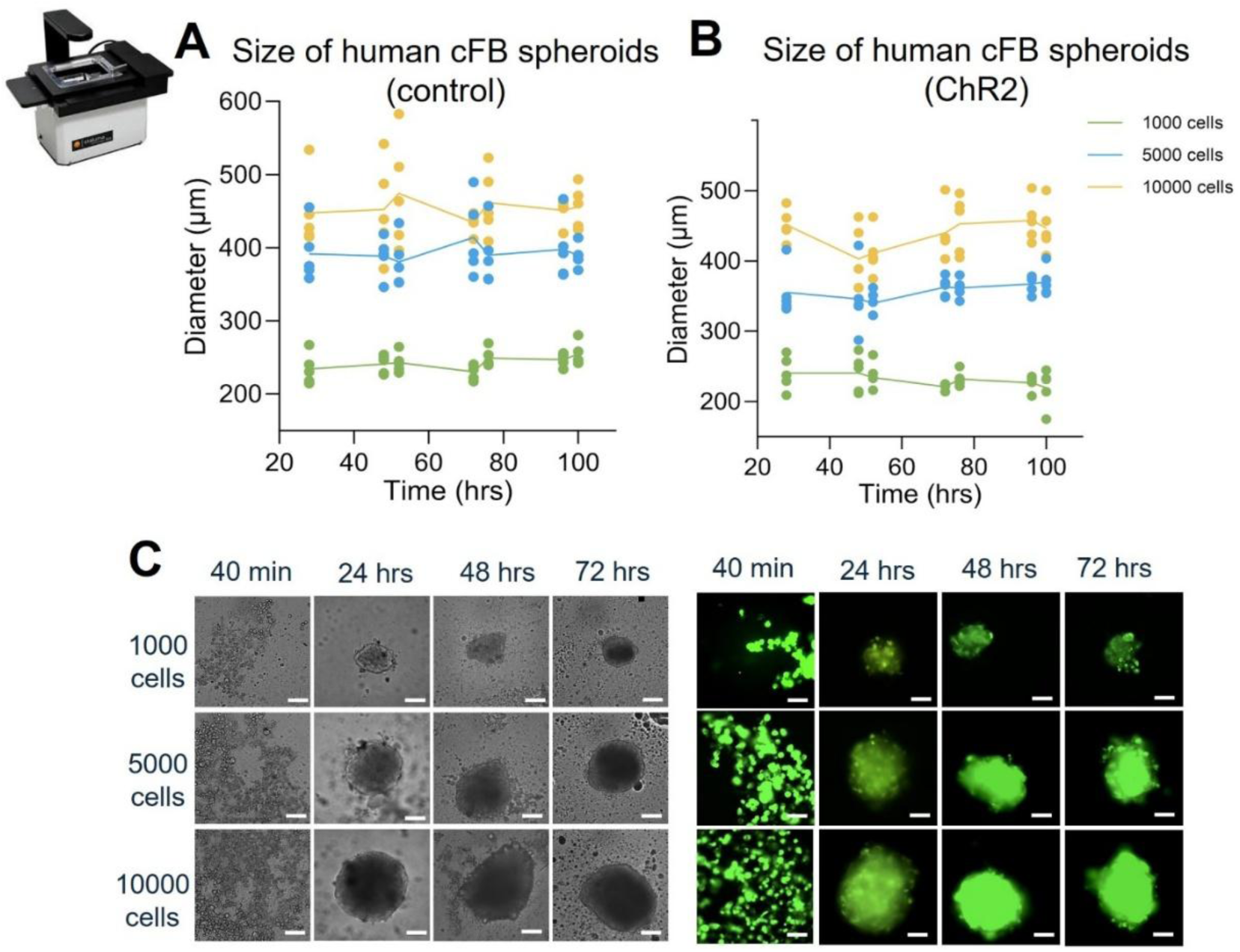
Automatic tracking of the assembly of optogenetically active human cardiac fibroblast spheroids: control and ChR2-modified cFB spheroids have similar growth and formation pattern. **A-B.** Diameter changes over time with control and Ad-ChR2 infected cFB spheroids at 1k, 5k, and 10k cells/spheroid (n=5 samples per size). **C.** Example cFB spheroid images for control (left) and +ChR2 (right) at different timepoints. Tracking was done within a standard incubator using an Etaluma microscope. Scale bars are 125 µm.

### Automated optogenetic control of cardiac assemblies using human cFB spheroids

Once we confirmed the utility of the Etaluma microscope for monitoring the cFB spheroid growth, we applied a similar approach for baseline characterization of the functional responsiveness (spontaneous rate) of hiPSC-CM cell spheroids when grown alone and when coupled with cFB cell spheroids. Spheroid formation for cardiomyocytes is much slower than cFB spheroid formation, happening over multiple days. Throughout that time, the iPSC-CMs begin clumping and beating together as one before a fully formed spheroid emerges. Not many previous studies have documented the long-term evolution of spontaneous rates of hiPSC-CM spheroids. To our knowledge, this is the first such application of the Etaluma platform for functional tracking of cardiac spheroids/organoids. Over the course of six days, we automatically tracked the beat rates of hiPSC-CM spheroids of different sizes (1k, 5k, and 10k cells) while growing inside a standard incubator. Across all sizes, there was a consistent decline in beat rate with time (culture medium was exchanged every two days). Furthermore, the larger spheroids were found to beat slower on average compared to the smaller spheroids, **Fig. 5A**. We also tracked the beat rates of size matched spheroid assemblies with CM and cFB spheroids, at the same three spheroid sizes, i.e. 1k CM – 1k cFB, 5k CM – 5k cFB, 10k CM – 10k cFB, over multiple days. Although the assembloids exhibited a substantially wider spread of beat rates compared to the CM spheroids alone, some interesting trends emerged. The smallest cFB - CM pairs (1k spheroids) tended to beat faster over time, rising to around 1.5 Hz on average, while the two larger spheroid pairs tended to decrease rate over time, trending towards quiescence, **Fig. 5B**.

**Figure 5.**
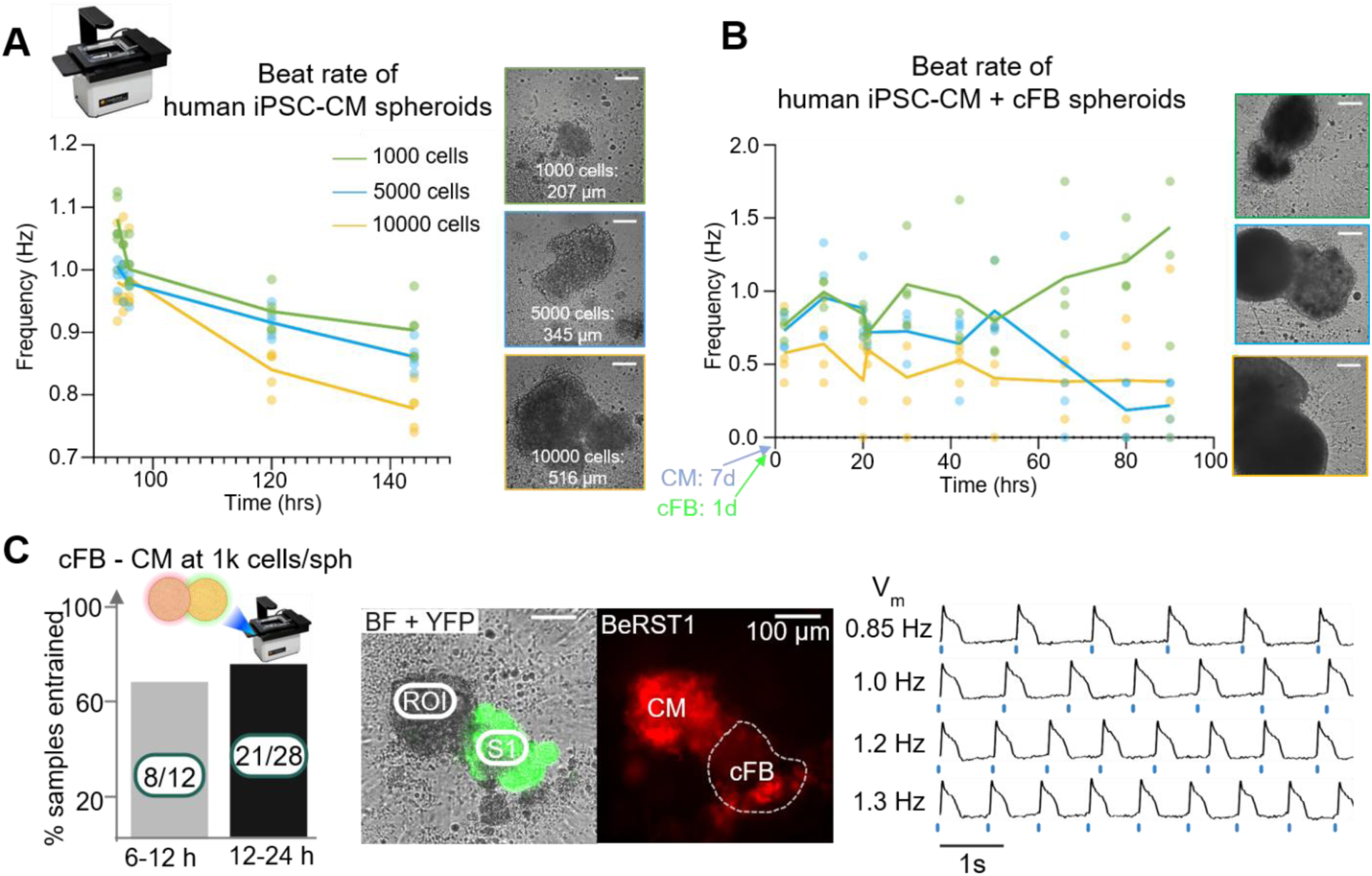
Automatic characterization of spheroid-spheroid assemblies and their actuation by light: cardiac fibroblast (cFB) spheroids can be used as effective optogenetic actuators for hiPSC-CM spheroids. **A.** Spontaneous rates over time for 1k, 5k, and 10k cell unmodified hiPSC-CM spheroids (n=5 per size). Example spheroid images are shown on the right of the graph. **B.** Spontaneous rates over time and example images for 1k, 5k, and 10k cell spheroid-spheroid assemblies using cFB and CM spheroids (n=5 per size), along with example images. Spheroid size and rate tracking was done using an Etaluma microscope in a standard incubator. CM spheroids were integrated 7 days post-plating with 1 day-old cFB spheroids. Scale bars in A-B are 125 µm. **C.** Left: Summary of sample entrainment for ChR2-cFB - CM spheroid pairings after 6-12 hours and 12-24 hours. Optogenetic probing of entrainment was done automatically with a modified Etaluma microscope inside a standard incubator. Middle: Example 1k ChR2-cFB - CM spheroid pairing, showing eYFP-ChR2 in green on the left and the voltage dye BeRST1 on the right, with respective recording and stimulation ROIs labeled. Example voltage traces from an entrained cFB - CM spheroid pair are shown on the right.

Next, we studied the entrainment of CM spheroids with ChR2-cFB spheroids (paired at 1000 cells each). We probed this both chronically inside the incubator and acutely outside the incubator using our all-optical OptoDyCE platform to record voltage responses upon optogenetic pacing. For the chronic probing over days, inside the incubator, for the first time, we modified the Etaluma microscope and the Lumascope software to automatically deliver optogenetic pacing and to track mechanical contractions. Entrainment was tracked over the course of 24 hours post-integration via the optogenetically-enabled Etaluma. The earliest entrainment for ChR2-cFBs – CM pairs was registered around 6 hours, longer compared to the ChR2-HEK. The cFB-actuated assemblies still performed well, with 66% (8/12) of the pairs being entrained for at least one frequency over multiple timepoints. This success rate rose to 75% (21/28) entrained after at least 12 hours of integration, **Fig. 5C**.

### Optogenetically-gated ultrasound stimulation and all-optical electrophysiology

We explored the extension of the TCU approach for cardiac pacing by other energies. Specifically, we were interested in cardiac pacing by ultrasound[22], directly in unmodified hiPSC-CM syncytia, in hiPSC-CM syncytia with sonogenetic enhancement and via the TCU approach using sonogenetically and optogenetically responsive cell spheroids. In this study, sonogenetic transduction was done by over-expressing human Piezo1[35, 36], a mechano-sensitive ion channel, using adenoviral delivery. Successful pacing was defined as 1:1 response rather than rate modulation by ultrasound. To perform these experiments, we constructed a specialized platform, where our all-optical OptoDyCE system (allowing for simultaneous optogenetic pacing while optically imaging voltage and calcium)[37, 38] was extended to accommodate also US stimulation from the top, **Fig. 6A**. A flat-face transducer was coupled using US gel to a film covering the solution-filled wells in a 96-well plate. All optical sensing and actuation were done through the glass bottom of the plate. Two function generators were used to synchronize the optogenetic and the sonogenetic stimulation with desired timings and pacing frequences. This allowed optogenetic gating and precise timing of the ultrasound pulses (select coupling intervals between the previous optogenetically triggered pulse and the US pulse) and immediate observation of both voltage and calcium responses. Example traces for three different US pulse intensities (0.5MHz, 1 to 6 W/cm^2^) are shown in **Fig. 6B** – following a train of optogenetically-triggered pulses, a single US pulse was delivered in this case, coupled at 0.5 sec from the delivery of the previous pulse. To estimate the actual pressure distribution within the HT well plate, we performed computational analysis, shown in **Fig. 6C**, showing peak acoustic pressures appearing at the cell/interface level. We also used an approximate method [39], converting the spatial-averaged peak intensity (I) to acoustic pressure (P), considering material density (cell culture medium), *ρ*, and speed of ultrasound waves in culture medium, *c*, **Fig. 6C**.

**Figure 6.**
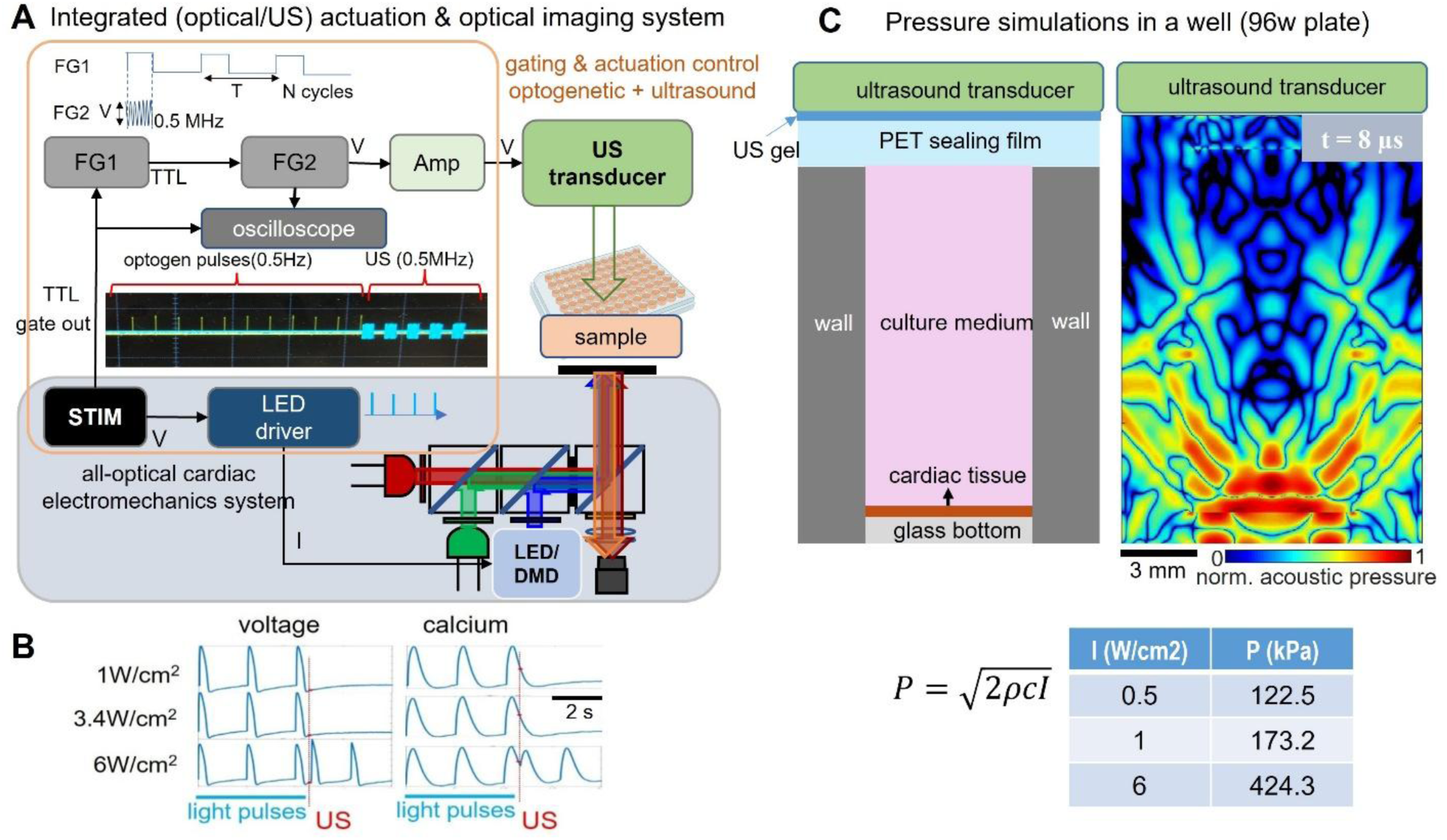
An integrated system allows multimodal actuation (optogenetic and US) and optical sensing of voltage and calcium. **A.** Integrated optical/US imaging and stimulation system schematic, with two function generators (FG1,2), amplifier, flat-face US transducer, LED driver, all-optical system for cardiac electromechanics and a 96-well plate with cells; the US transducer was coupled to the samples via gel on top of a thin film over the solution in the wells. **B.** Representative voltage and calcium traces in response to optogenetic pacing (0.5 Hz) followed by a delivery of an ultrasound pulse (coupling interval 0.5 s, duration 0.5 s) with different acoustic intensity (W/cm^2^) in hiPSC-CMs. **C.** Acoustic propagation and pressure distribution modeling within a 96-well plate: schematic with modeled materials and a snapshot of acoustic pressure distribution at an 8 µs propagation time, scale bar for the schematic and simulation is 3mm, normalized acoustic pressure shown. Below: simplified conversion of spatial-averaged peak intensity (I) to acoustic pressure (P), considering material density (cell culture medium), *ρ*, and speed of ultrasound waves in the culture medium, *c*, as per [39].

### Human Piezo1 as a sonogenetic tool: endogenous cardiac expression and viral delivery

Piezo1 has been previously suggested as a contributor to the responses to US stimulation in excitable cells[36, 40]. Its expression in the heart, and specifically in cardiomyocytes, is not very well characterized. We used qPCR to quantify the endogenous Piezo1 expression in our human iPSC-CMs and in adult human cFBs, **Fig. 7A**. We found that the cFBs had substantially higher native Piezo1 expression, while the hiPSC-CMs had minimal levels. Adenoviral delivery of human Piezo1 at different multiplicity of infection (MOI) substantially boosted expression in both cell types, especially in the hiPSC-CMs. Furthermore, we used RNAseq to compare the Piezo1 expression from hiPSC-CMs and the adult human ventricle (using the GTEx database), after similar normalization of the transcriptome, **Fig. 7B**. Compared to other key cardiac ion channel genes (sodium and calcium), the expression of Piezo1 is lower, but follows a similar relative pattern in the adult heart and in the hiPSC-CMs. Finally, we optimized the generation of human HEK cell spheroids co-expressing both ChR2 and Piezo1 for combined optogenetic and sonogenetic actuation, see **Fig. 7C**. The viability, growth and performance of these combined action spheroids remained unaltered with the optimized expression.

**Figure 7.**
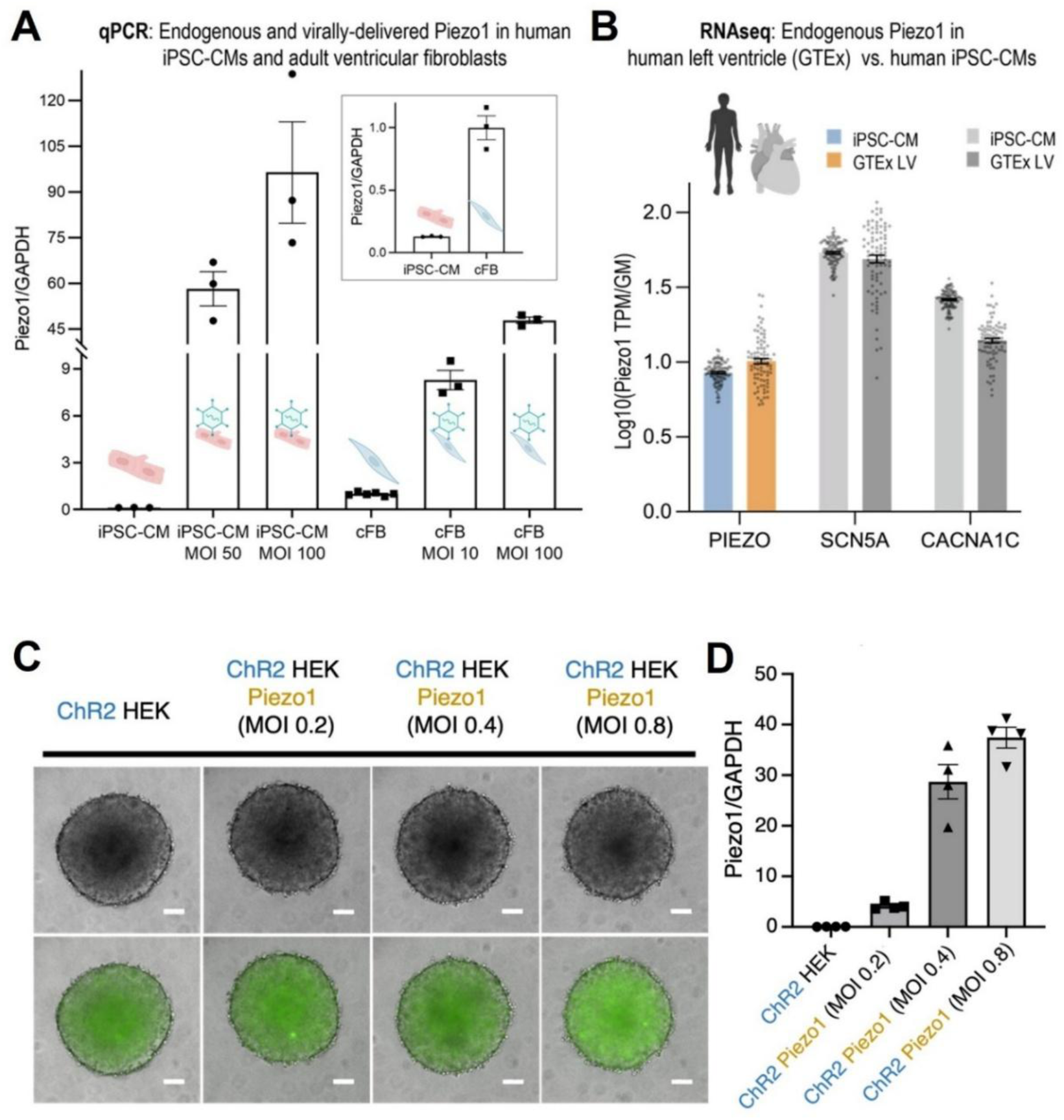
Endogenous and exogeneous expression of Piezo1 in iPSC-CMs and cFBs and generation of optogenetically and sonogenetically active cell spheroids. **A.** Relative Piezo1 mRNA quantification of endogenous and virally transduced Piezo1 in human hiPSC-CMs and adult ventricular fibroblasts. **B.** Comparison of relative Piezo1, SCN5A, and CACNA1C mRNA levels in hiPSC-CMs and human left ventricle tissue. **C.** 10K spheroid assembly of ChR2-eYFP only, and ChR2 and Piezo1 expressing HEK cells at respective MOIs. **D.** mRNA quantification of Piezo1 at the respective relative MOIs. Scale bars are 100 μm.

### Combined optogenetic and sonogenetic actuation for cardiac pacing

To simulate the optogenetically- and sonogenetically-triggered cardiac responses, we used an electro-mechano-metabolic computational model of human iPSC-CM[41], in which we introduced our previously published ChR2 model[42] and a modified version of our simplified Piezo1 model[43], **Fig. 8A**. Optogenetic pacing with short (10 ms) pulses at 0.15 mW/mm² (similar values to the ones we used experimentally) yields action potentials that are similar in morphology to electrically triggered ones, as shown previously[44]. Mechanical (ultrasound) stimulation was implemented by applying effective stress of 0.336kPa on the Piezo1 gate – a stress that was 3.4 times the peak native cell stress from regular contractions and twice the lowest tested amplitude that triggered an action potential (0.168 kPa), for a cell with a selected whole-cell conductance of 9 nS for functional Piezo1 channels (having single channel conductance of approximately 29pS [45]). In this study, we did not attempt to link the acoustic intensity to actual membrane stress. Note the very similar but slightly shorter action potential triggered via 0.5 s of mechanical stimulation through the Piezo1 channels. Both stimuli initiate excitation through a transient inward current; ChR2 and Piezo1 also carry a very small outward current during part of the action potential. Based on the relatively quick inactivation of both ChR2 and Piezo1 upon triggering a cardiac action potential, future light and ultrasound protocols do not need to use stimuli longer than 30 ms. Overall, stimulation by light and by ultrasound could be shown computationally for reasonable stimulation parameters (irradiance and mechanical pressure). Further modeling is needed to link the particular acoustic pressures to mechanical stress exerted on the membrane.

**Figure 8.**
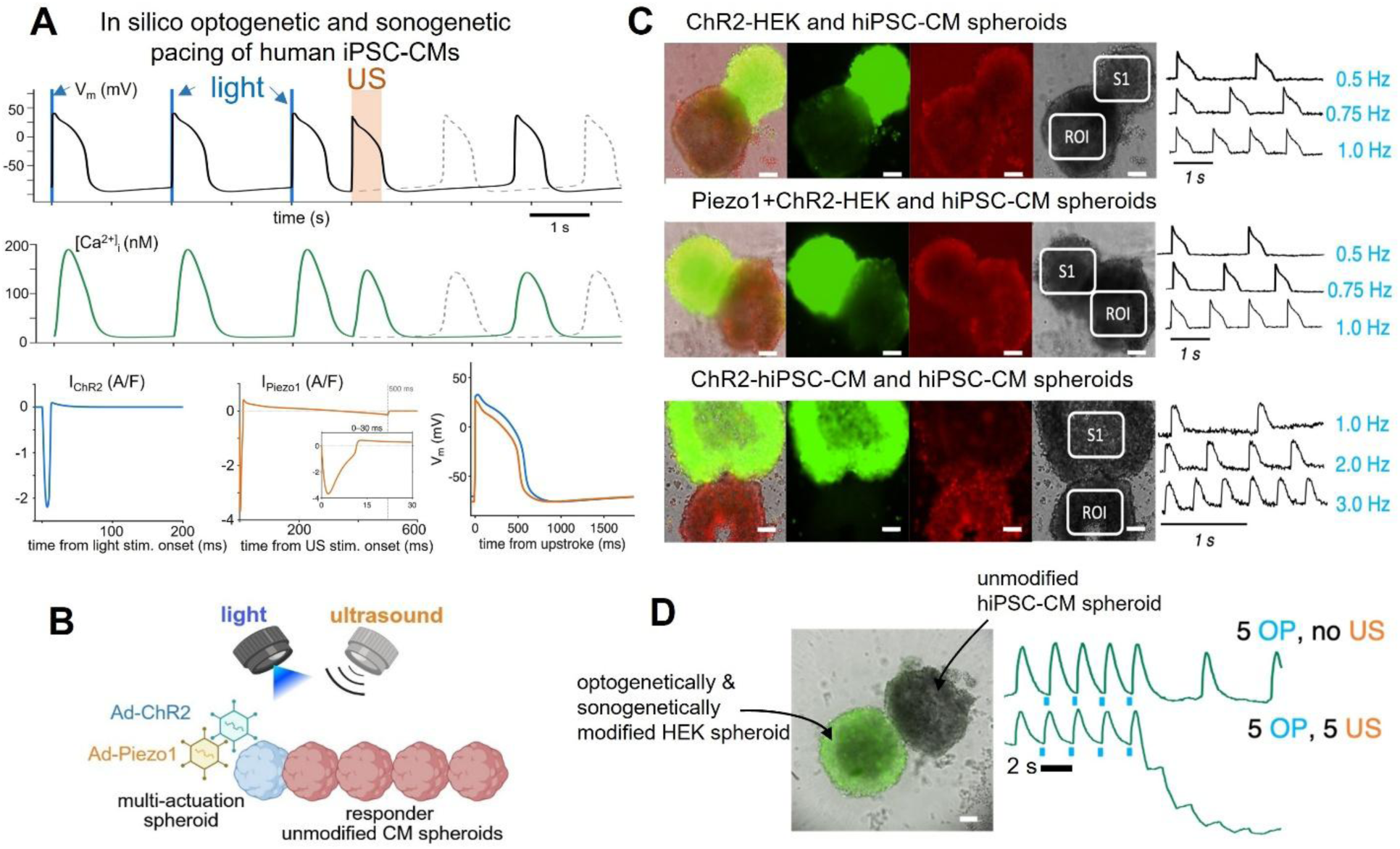
In silico optogenetic and sonogenetic pacing of hiPSC-CMs and experiments using hybrid spheroid pair assemblies. **A**. Simulated response of a single hiPSC-CM at 31 °C to three optical pulses (10 ms, 0.15 mW/mm², 0.5 Hz), followed 1 s after the third pulse by a 0.5 s mechanical stress pulse (0.336 kPa effective stress at the Piezo1 gate). Shown are action potentials, calcium transients and ChR2 and Piezo1 currents; the dashed gray trace is the no-mechanical-stimulus control. Optical and mechanical action potentials are aligned at their upstrokes for comparison. The mechanically triggered AP is slightly shorter than the optogenetically-triggered AP. **B**. Envisioned hybrid spheroid assemblies responsive to optogenetic and sonogenetic pacing. **C.** Examples of responses to optogenetic pacing in spheroid-spheroid assemblies, as shown (10k cell spheroids). D. An example of optogenetic gating of ultrasound application: pacing by light followed by zero or five ultrasound pulses in a cell spheroid pair (Piezo1+ChR2-HEK and hiPSC-CMs, 10k). Scale bars are 100 μm.

Experimentally, we combined optogenetically and sonogenetically responsive cell spheroids with unmodified hiPSC-CM spheroids, **Fig. 8B-C**, to confirm the preservation of their ability to pace optically (**Fig. 8C**) and to test their ability to confer responsiveness to ultrasound, **Fig. 8D**. Note the 1:1 response to a pacing sequence of multiple optogenetic pulses and multiple US pulses.

Ultimately, we performed a comprehensive study evaluating the responsiveness of various configurations for optogenetically-gated ultrasound pacing of hiPSC-CMs, **Fig. 9**. Our previous work has established that tailored optogenetic modifications do not alter electromechanical responses in cardiomyocytes[38]. We did not observe overt changes in Piezo1-modified hiPSC-CMs in this study, nor in the spheroids with these sonogenetic modifications. In some cases, we used a pharmacological activation of Piezo1 with a specific agonist – Yoda1 (at 10 µM). This was a combinatorial study of 16 conditions, where in each case optogenetic pacing was followed by testing the responsiveness to a sequence of mechanical stimuli: +/-US, +/-Piezo1 expression, +/Yoda1 activation of Piezo1, and +/-TCU spheroid-mediated action. We classified the responses to five delivered US pulses as no response, intermediate (meaning change in spontaneous rate) and 1:1 response pacing.

**Figure 9.**
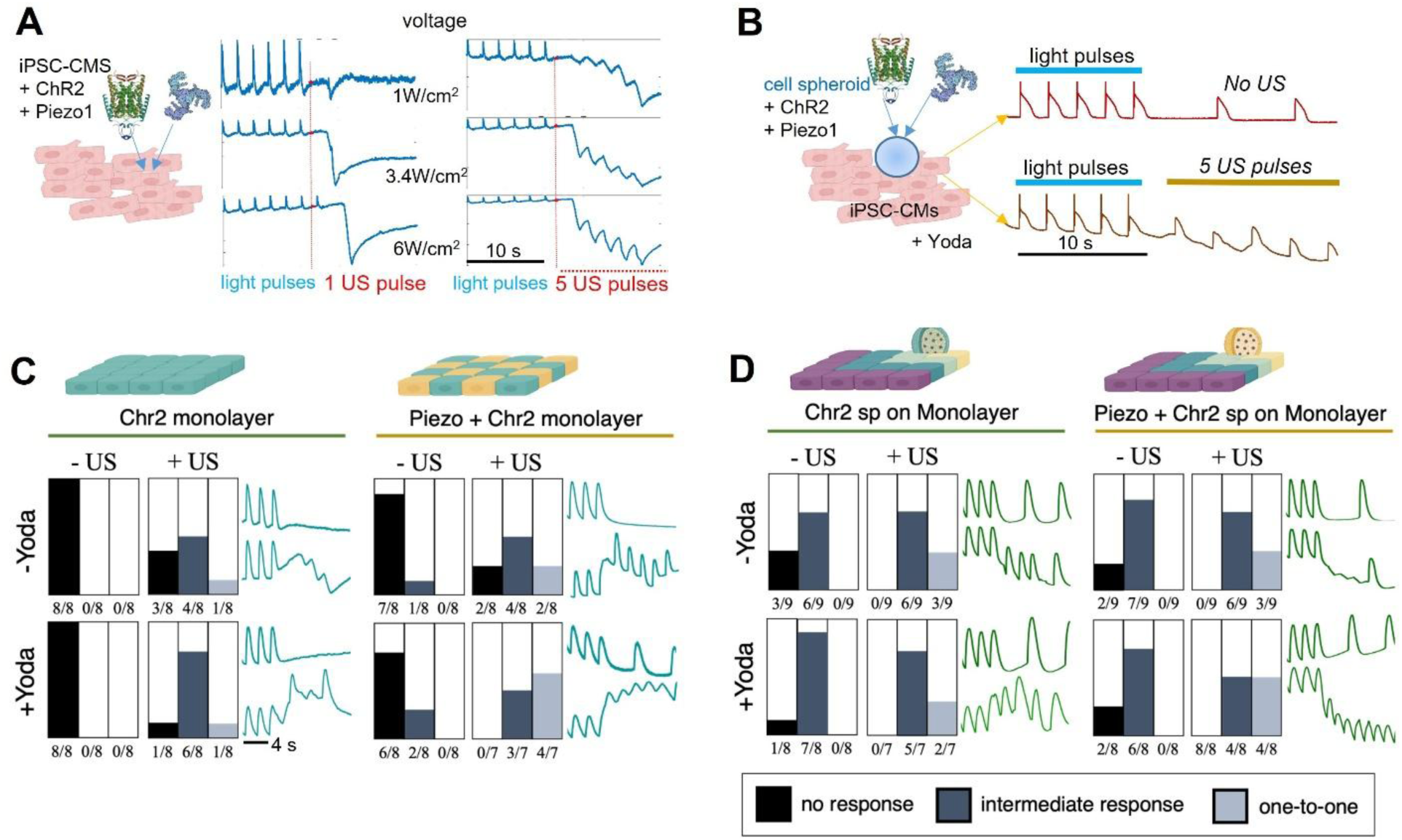
Ultrasound pacing of human iPSC-CMs with optogenetic gating in various configurations. **A-B.** Schematic and example voltage traces from optogenetic and sonogenetic pacing of modified iPSC-CM monolayers and unmodified iPSC-CM monolayers via optogenetically and sonogenetically-enhanced cell spheroids, respectively. **C-D.** Quantification of responses of iPSC-CM monolayers to light (0.15 mW/mm^2^) and ultrasound (1 W/cm^2^) for the two configurations shown in A and B, with and without Yoda1 agonist (10 µM) of Piezo1, n=8 samples per condition (16 different conditions). Three types of responses were documented: no response, intermediate (change in rate) response and one-to-one pacing. Example traces are shown to the side for -/+ US without Yoda1 and -/+ US with Yoda1 activation for the respective cases.

In all tested conditions, the absence of US never resulted in 1:1 pacing. In all studied groups, +US induced increase in the combination of intermediate and 1:1 responses compared to -US. In hiPSC-CM syncytia without sonogenetic modification, Yoda1 application at baseline (-US) did not induce any intermediate or 1:1 effects (**Fig. 9C**), consistent with the relatively low gene expression of native Piezo1. However, US application in such unmodified hiPSC-CMs did induce some intermediate rate changes and minimal 1:1 responses (1/8 samples). Sonogenetically modified syncytia exhibited enhanced responsiveness to US, especially with Yoda1 activation (1:1 responses in 4/7 samples).

Of most interest to us was the testing of the TCU approach to mediate ultrasound effects on pristine (unmodified) hiPSC-CM syncytia (**Fig. 9D**). The deployment of dedicated biological actuators (optogenetic/sonogenetic spheroids) increased the non-specific (not US-related) change in rate at baseline compared to hiPSC-CM syncytia without cell spheroids. Because of movement upon US application, the optical signals had a shifting baseline. However, the TCU approach worked as expected to enhance the specific (intermediate + 1:1) responsiveness to US, in a manner similar to direct sonogenetic modification of the cardiac monolayers. This is in line with our experience with optogenetic actuation – direct and via TCU. Yoda1 activation increased the 1:1 responsiveness to US in the group with sonogenetically-active cell spheroids. While none of the tested conditions resulted in 100% 1:1 US pacing at this intensity (1W/cm2), we were able to corroborate many factors leading to enhanced sonogenetic actuation – Piezo1 overexpression, Yoda1 activation and TCU-mediated responsiveness using dedicated biological actuators.

## DISCUSSION

In this work, we demonstrate that both contactless optical and ultrasound-mediated actuation of unmodified cardiac tissue can be achieved after short-term integration of designer cell spheroids, without requiring genetic modification of the target myocardium. These designer spheroids for multi-modal actuation can be magnetically guided to a site of deployment and then engaged by external light or ultrasound source. In short, we report that the TCU concept is applicable across different energy modalities and furthermore, we quantify the factors that help such indirect stimulation. This approach, leveraging human cells, potentially patient-derived, that do not need to be excitable, has a multitude of potential applications in cardiac research, both *in vitro* and *in vivo*.

For example, the biological actuators can be directly used as a reagent in high-throughput cardiotoxicity screening, for scalable point-pacing without contact electrodes and without genetic modification of the assayed cells/tissue, as shown recently in all-optical electrophysiology platforms[46]. We have shown previously that such cell spheroids can be cryopreserved and then thawed and used as live bioactuators without loss of performance[26]. Furthermore, considering the growing interest in cardiac assembloids and organoids [30, 47] as human-based model systems for characterizing human cardiac function in health and disease, insights from our study can be used to empower these efforts by adding the ability for control during organogenesis by various energy modalities using modular assembly including the described here bioactuators. Overall, the described technology is in line with initiatives to develop new approach methodologies (NAMs) to enhance human-based alternatives to animal studies in preclinical research[48].

For *in vivo* applications, optogenetic pacing faces multiple translational challenges[13, 14], where the gene modification/therapy aspects constitute a major hurdle. Delivery of specialized biological actuators formed out of patient’s cells, similar to the ones characterized here, may alleviate these concerns, as it avoids uncertainties of non-specific targeting of viral vectors *in vivo*. Combined with newer very-high conductance opsins with red-shifted spectral profile[15, 16], such magnetically-guided bioactuators can exert rate control from well-defined locations in the heart, triggered by external or subcutaneously placed LEDs. The minimally invasive delivery of cell spheroids has been demonstrated in recent heart regeneration clinical trials[27–29, 49]. In fact, a potential clinical application would be to augment such heart regeneration efforts by deliberately incorporating optogenetic or sonogenetic biological actuators for phase resetting and temporary control of rate during the transient period of integration to avoid engraftment arrhythmias[3]. This application would only require engagement of the biological actuators over the initial few weeks. Such engagement by light or ultrasound, with real time feedback based on pre-derived patient-specific phase resetting curve[3] and the patient’s ECG is ideally suited to seamlessly synchronize the competing pacemakers (the native sinus node and the newly introduced engineered heart tissue), after which external pacing may no longer be needed. With optimal choice of the cells used in these biological actuators (ideally, patient-derived), one may potentially arrive at a superior solution for transient pacing compared to the man-made bioresorbable pacemakers, where the byproducts may irritate the underlying tissue during biodegradation.

Sonogenetic cardiac pacing or probing of activation patterns by US *in vivo* is motivated by the deeper penetration depth of ultrasound stimuli compared to optogenetic stimulation even with the furthest near-infrared responsive opsins. Newer developments of miniaturized wearable ultrasound transducers suggest that such devices are a possibility[24, 25, 50, 51]. Ultrasound-based probing for device placement for cardiac resynchronization therapy (CRT) has already been used in humans [5, 52]. Sonogenetics via distributed cell delivery locations can be potentially helpful in these efforts. In this study, we focused on sonogenetics with a particular mechanosensitive ion channel, Piezo1, that has been implicated in responses to ultrasound[35, 40, 53, 54]. However, there are a variety of other options, some of which have been explored in other studies, including native mammalian calcium-conducting ion channels TRPA1[55], two-pore-domain potassium ion channels (K2P), e.g. TREK-1, TREK-2, TRAAK [22] or exogenous bacterially-derived mutant large-conductance ion channels MscL-G22S [24]. Any of these sonogenetic enhancers can be combined with our TCU approach. The dedicated biological sonogenetic actuators (cell spheroids) can mediate/amplify the ultrasound action upon integration with unmodified native tissue, and represent an attractive opportunity for well localized pacing, similar to TCU’s applicability in optogenetics. Sustained mechanical pacing is more challenging than sustained electrical or optogenetic pacing. Combined responsiveness to light and US offers additional flexibility to exploit synergistic modulation of membrane potential. Computational modeling, as exemplified here, can be further developed to offer a digital twin platform for optimizing new technologies for cardiac actuation and guiding the choice of actuators and stimulation protocols that can perform well for pacing or anti-arrhythmic action at the lowest energy requirements. Our current models elucidate the electromechanical responses to light or ultrasound, as mediated by the triggered ionic currents through the light- and mechano-sensitive ion channels. Future developments are needed to translate the irradiance at the light source to irradiance experienced at the cell membrane using photon migration modeling; similarly, future modeling needs to link the ultrasound acoustic pressure to mechanical stress exerted on the cell membrane for immediate assessment of optimal stimulation parameters. Ultimately, *in vivo* testing will be needed to confirm the practical utility of the approach to cardiac pacing.

Overall, this study explored the fundamental possibility to develop new alternative methods for cardiac pacing and rhythm control. In the process, we also developed some new platforms and computational tools for comprehensive functional analysis aiding new developments in this area using human/patient-derived cells.

## METHODS

### Cell culture: human iPSC-CMs, human cardiac fibroblasts (cFB) and human embryonic kidney cells HEK293T

For monolayer growth, iCell Cardiomyocytes^2^ (Cat. C1016, Donor 01434, Fujifilm/Cellular Dynamics), were cultured on fibronectin-coated (50μg/mL) glass-bottom 96-well plates at 50,000 cells per well and maintained according to manufacturer’s protocol at 37°C and 5% CO_2_. For spheroid formation or other formats, see below.

Adult human ventricular cardiac fibroblasts (cFBs, Cell Applications, Inc. 307V-75a) were thawed and cultured according to manufacturer’s instructions. Cells were used within five passages.

We have formerly developed[19] an immortalized 293T-HEK cell line expressing the optogenetic actuator, Channelrhodopsin-2, ChR2, tagged with yellow fluorescent protein (YFP) using the Addgene construct pcDNA3.1/hChR2(H134R)-EYFP, provided by Karl Deisseroth. ChR2-HEK cells were thawed, plated, and cultured in Dulbecco’s Modified Eagle Medium (DMEM, ATCC) with 10% fetal bovine serum (FBS), 1% 200 mM L-Glutamine, and 1% penicillin-streptomycin until confluent.

### Gene delivery of optogenetic and sonogenetic actuators

Optogenetic responsiveness in the hiPSC-CMs and the cFBs was achieved using an adenoviral delivery of the opsin, Ad-CMV-hChR2(H134R)-eYFP (Vector Biolabs), at multiplicity of infection, MOI of 50 and 200, respectively, following previously validated protocols[34, 56]. Expression was present after 24 to 48 hours.

Sonogenetic responsiveness (in hiPSC-CMs, cFBs and HEK 293T) was enhanced using a delivery of the human Piezo1 mechanosensitive ion channel via an adenoviral vector Ad-h-Piezo1 (ADV-231912, Vector Biolabs) at MOI 50 or as indicated in the results. Expression was confirmed with qPCR after 48 hours. In some experiments with ultrasound, we used the Piezo1 agonist, Yoda1 (Cat. 1558, Sigma-Aldrich), diluted to 10μM and applied acutely to the cells.

Co-expression of optogenetic and sonogenetic actuators was optimized by varying the sequence of administration and the dose, as reported. In all cases, the viability/health of the cells was not affected at the doses used.

### Cell spheroid formation

For spheroid assembly, cFB and ChR2-HEK were rinsed with phosphate-buffered solution (PBS) and detached using 0.05% trypsin-disodium ethylenediaminetetraacetic acid (EDTA) in Hank’s Balanced Salt Solution (HBSS). The cells were centrifuged and resuspended at the desired plating density before seeding. In contrast, the hiPSC-CMs were used for spheroid formation directly after thawing. Because hiPSC-CM spheroids were found to be healthy and beating up to three weeks after seeding, they were used 6-21 days post-thawing. Most of the HEK 293T and cFB spheroids were used within 2-3 days after formation.

Two methods of spheroid formation were applied. One method used ultra-low-adhesion 96-well spheroid microplates by Corning (Catalog #CLS4920, Millipore Sigma) to form one spheroid per well at the desired density. A second method for mass production used EZsphere 35mm dish (#4000-900, Iwaki) to form 2700 cell spheroids (1000 cells each) at once. Cell spheroids were maintained by 50% medium exchange every two days.

### In-incubator long-term tracking of cell spheroid assembly and spontaneous rate

Automated long-term (up to 8 days) tracking of cell spheroid formation and activity, as well as spheroid pairing was done using Etaluma LS850 fluorescence microscope with custom-added capability for optogenetic pacing in a standard incubator. Static images and videos were acquired at 20X magnification. Optical pacing used 488 nm LED, 10 ms light pulses at varying pacing frequencies. Videos were recorded at 20-25 fps in brightfield or in the red channel (612-680nm), if stained with a NIR voltage-sensitive dye. The acquired data were used to quantify cell spheroid size over time post assembly and to extract spontaneous or entrained beat rate, using Matlab scripts.

### Magnetic nanoparticle incorporation in cell spheroids and spheroid positioning

For responsiveness to magnetic field, cells were exposed to media containing magnetic nanoparticles, mNPs, at varying dilutions from the manufacturer’s product. We used primarily Greiner mNPs (NanoShuttle, #657841; Greiner Bio-One) at 1:10 dilution. These 50nm mNPs adhere stably to the cell membrane or get endocytosed, as previously described[57–59]. Additional mNPs tested here were Takara mNPs (Lenti-X Accelerator, #631257; Takara Bio) at 1:200 dilution and Micromod mNPs (nanomag-CLD-redF, #23-01-102; Partikeltechnologie GmbH) at 1:50 dilution. This was done mostly while the cells were in a monolayer format, after overnight incubation with the mNPs, the cells were lifted and seeded as spheroids. In the case of hiPSC-CM spheroids, mNPs were introduced in the early hours of spheroid formation. At the reported concentrations, all cell spheroids became responsive to external guidance without effects on cell health, and could be easily pulled down by a magnetic plate under the 96well plate or localized to the center vs. edge of the dish.

### Deposition of spheroids onto cardiomyocyte syncytia for TCU studies

Pre-formed optogenetic/sonogenetic spheroids (HEK 293T or cFB), with or without mNPs, were deposited on unmodified hiPSC-CM monolayers grown in glass-bottom 96well plates using a cut-tip P200 pipette, similar to our previous study [26]. A subset of studies were done with hiPSC-CMs grown on quasi-3D electrospun scaffolds (700 nm polycaprolactone or PCL fibers) in special 96well plates from Nanofiber Solutions (SKU 9601, 9602), as in prior work[46]. For cases where the cell spheroids had mNPs, the samples were placed on a magnetic plate for 10-15 min or spheroids were specifically magnetically guided to the center or the periphery of the well and then further incubated for 1-2 hours before experiments. The cardiac monolayers were pre-labeled with voltage/calcium dyes before spheroid integration, and then functional tests were carried out in Opti-MEM medium (#11058-021, Gibco).

### Spheroid-spheroid pairing for TCU studies

Hybrid pairs of cell spheroids were formed by co-introducing the already formed spheroids into ultra-low-attachment 96well or 384well plates. One of the spheroids in a pair was with enhanced actuation to light (and US) and the other was a unmodified hiPSC-CM spheroid, pre-labeled with voltage/calcium dye. Typically, spheroids were size-matched, however a select number of experiments were conducted with deliberately mis-matched in size cell spheroids. Probing of “drivability” (or entrainment by light or US) was done within 1 to 2 hours after pair co-introduction.

### qRT-PCR analysis

Cells were harvested for total RNA and quantified using the POWER SYBR^TM^ Cells-To-C_T_ Kit (Cat. 4402955, ThermoFisher Scientific) following the manufacturer’s protocol. qPCR analysis was performed on the QuantStudio 3 Real-Time PCR System (ThermoFisher Scientific) to detect each target gene and analyzed using QuantStudio^TM^ (ThermoFisher). Primers used for mRNA detection of human *Piezo1* were Fw_GCCGAGAGACAGAGAAGAAATAC and Rv_GCGATGAGGAAGAGGATGATG; and for detection of *GAPDH* were Fw_GGAGCGAGATCCCTCCAAAAT and Rv_GGCTGTTGTCATACTTCTCATGG. All samples were normalized to housekeeping gene GAPDH and results were expressed as the relative mRNA normalized to control using the standard ΔΔCt method, as described previously[60].

### RNAseq analysis of Piezo1 expression in the adult human heart (GTEx database) and in hiPSC-CMs

We checked for endogenous Piezo1 expression in the adult human heart (left ventricle) and human iPSC-CMs using RNAseq. Data from the GTEx database, version 8 [61] and our own iPSC-CM data were normalized the same way and gene expression was compared, similar to the methods in [62, 63].

### Immunocytochemistry

Immunocytochemistry was performed to visualize spheroids on the 3D scaffolds. ChR2-HEK spheroids were identified by their eYFP tag. iPSC-CM spheroids were labeled for α-actinin with a monoclonal antibody (Catalog #A7811, Millipore Sigma) and for nuclei with Hoechst (Catalog #H3570, Thermo Fisher Scientific). Fixed cells were rinsed with PBS, permeabilized with 0.2% Triton X-100 in 5% FBS, labeled via primary and secondary antibody staining, and imaged using confocal microscopy.

### All-optical electrophysiology probing for entrainment

Measurements of functional responses to pacing by light for spheroid-monolayer/electrospun matrix and spheroid-spheroid pairings, were conducted using our OptoDyCE system built around an inverted Nikon Eclipse Ti2 microscope at 20× magnification[37, 38]. Optical stimulation was delivered via a digital micromirror device (DMD; Polygon 4000, Mightex, Toronto, ON, Canada) using 470 nm blue light pulses of 10 ms duration, <0.15 mW/mm² intensity, and varying frequencies to engage the modified spheroids. Membrane voltage and calcium transients in the responding hiPSC-CMs were monitored with the BeRST1 dye (1 µM) under 660 nm excitation and Rhod-4 (10 µM) excited with 535nm, respectively, using temporal multiplexing. The recording camera was iXon Ultra 897 EMCCD camera (Andor Technology Ltd., Belfast, United Kingdom) at 200 fps. Regions of interest for DMD-based optical stimulation and optical voltage recording were manually selected to restrict stimulation to the optogenetically-modified spheroid and sensing to the iPSC-CM syncytium or spheroid. For each sample, these regions were kept consistent between recordings. Signals were filtered and analyzed using automated custom software to quantify relevant parameters. Entrainment was marked with 1:1 following of at least two distinct pacing frequencies.

### Integrated system for ultrasound/optogenetic stimulation and optical imaging

The integrated ultrasound/light system is shown in **Fig. 6A**. A 0.5-MHz immersion ultrasound transducer (17-0012-P-SU, Olympus) was utilized to deliver ultrasound stimulation. The transducer was integrated into the all-optical imaging system using a customized mount and a 3-D translational stage. For acoustic alignment, the microscope’s transmitted illumination light served as a spatial reference to center the transducer over the target well containing the cardiomyocytes. Following alignment, ultrasound coupling gel was applied between the PET sealing film on top of the 96-well plate and the transducer surface, with air bubbles carefully removed to guarantee efficient acoustic transmission. The timing synchronization and signal generation for the combined ultrasound and optogenetic stimulation were coordinately controlled by a field stimulator (MyoPacer EP, IonOptix), two waveform function generators (DG1032Z, Rigol), FG1,2, and an RF power amplifier (240 L, E & I). Upon initiating the multi-channel optical imaging to capture voltage and calcium responses, the field stimulator first applied a 20-sec optogenetic pacing sequence (0.5 Hz, 11 blue light pulses). Following the final optical pulse, the first waveform generator introduced a coupling interval before triggering the second generator to output one or five bursts of a 0.5-MHz, ultrasound excitation signal (up to 1 sec in duration) at specific voltage amplitudes (*V*). The RF amplifier (50-dB gain) drove the piezoelectric transducer to target acoustic intensities (*I*), calibrated to a baseline voltage (*I* ∝ *V*^2^)[39, 64]. Throughout the combined stimulation, the optical imaging system continuously recorded the voltage/calcium transient signals at 200 fps, as described above.

### Modeling of acoustic propagation and pressure distribution in a 96 well plate

To evaluate the acoustic wave propagation and pressure field distribution within a single well, a computational model was developed using COMSOL Multiphysics software (version 5.6, COMSOL Inc), **Fig. 6C**. The simulation geometry replicated a single test well (6.5 mm in diameter) coupled with a 0.5-MHz unfocused ultrasound transducer (23 mm in diameter). Along the acoustic propagation axis, the model incorporated the culture medium (11-mm height), the target cardiac tissue layer (20-μm thickness), and a Pyrex glass bottom (1-mm thickness), radially bounded by polystyrene walls. Empirical acoustic properties, specifically the speed of sound (distinguishing longitudinal, *c_L_*, and shear, *c_s_*, wave velocities where applicable), density (*ρ*), and acoustic attenuation coefficient (μ*_a_*), were assigned to each material domain: culture medium (*c_L_*: 1496 m/s, 998 kg/m³, 0.0022 dB/cm), cardiac tissue (*c_L_*: 1540 m/s, 1081 kg/m³, 0.52 dB/cm), Pyrex glass bottom (*c_L_*: 5640 m/s, *c_s_*: 3280 m/s, 2320 kg/m³, ∼1 dB/cm), and polystyrene walls and top film layer (*c_L_*: 2350 m/s, *c_s_*: 1120 m/s, 1060 kg/m³, 3.6 dB/cm). A 0.5-MHz continuous wave excitation was applied at the transducer boundary.

The spatial-averaged acoustic pressure was defined based on the theoretical intensity-to-pressure conversion 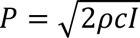,[39], where an intensity of 1 W/cm^2^ corresponded to an acoustic pressure of 173.2 kPa. The simulation mapped the spatial distribution of the normalized acoustic pressure field, verifying the acoustic stimulation delivered to the cardiomyocytes layer. According to [39], the acoustic pressure (*P*) is calculated by:

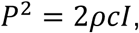

where *ρ* is the density of cell culture medium, *c* is the speed of ultrasound waves in the culture medium, and *I* is the ultrasound intensity. For simple calculation, *ρ* and *c* can refer to the corresponding values of them in water (10^3^ kg/m^3^, 1500 m/s; independent of ultrasound frequency). Then we have the following pressure calculation results, **Fig. 6D**.

For example, for *I* = 1 *W*⁄cm^2^ = 10^4^ *W*⁄*m*^2^, we obtain

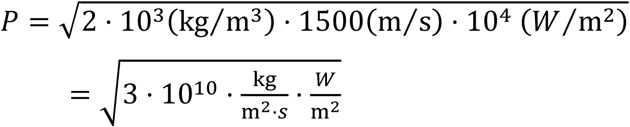

As 1 *W* = 1 *N* ⋅ m⁄*s* and 1 *N* = 1 kg ⋅ m⁄*s*^2^, the calculation is rewritten as

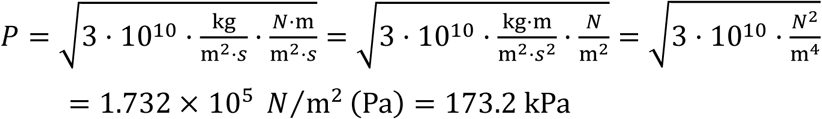

### Computational modeling of optogenetic and sonogenetic stimulation of human iPSC-CMs with ChR2 and Piezo1

For the simulations we used our published model of human iPSC-CMs - hiMCES2026[41], with inserted ChR2 model of Williams et al.[42] and a modified Piezo1 model based on our published CinC2023 formulation[43]. Simulations were performed at 31 °C; experiments were conducted at room temperature. Intrinsic host parameters were retained; ChR2 and Piezo1 currents were added to the model. For the Piezo1 description, the voltage-sensing gate from [43] was replaced by an availability gate, and three kinetic parameters were fitted to published channel recordings[65–67]. Effective functional Piezo1 channel number per cell was represented by a whole-cell conductance of 9 nS, corresponding to approximately 310 channels using a nominal unitary conductance of 29.1 pS[45]. Native cellular tension reached the gate through a first-order filter with a 0.15 s time constant; the imposed mechanical stimulus acted directly on the gate.

In the simulations, cells were conditioned by 0.5 Hz optogenetic pacing (10 ms at 0.15 mW/mm²). The responses to three optical pulses are shown, followed 1 s later by one 0.5 s mechanical stress pulse of 0.336 kPa representing the US stimulus. No acoustic propagation or conversion from acoustic intensity to gate stress was modeled. Equations were integrated with ode15s in MATLAB R2026a.

The Piezo1 current is represented with a Hodgkin-Huxley-type formalism:

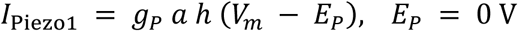

The activation gate, *a,* and availability, *h*, are dimensionless.

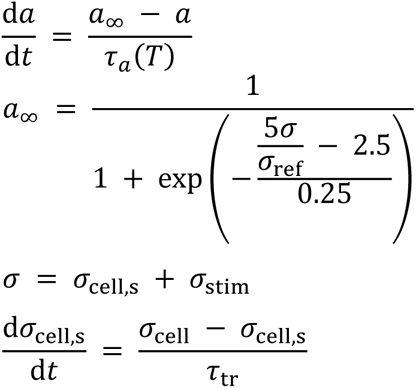

*I*_Piezo1_ is current per capacitance (A/F), *g*_P_ maximal conductance per capacitance (S/F), *V*_m_ membrane voltage (V), and *E*_P_ reversal potential (V). *a*_∞_ is steady-state activation and *τ*_a_ the activation/deactivation time constant. Stress terms (kPa) are σ (total sensed stress), *σ*_cell,s_ (filtered native stress), *σ*_stim_ (applied stress), and *σ*_ref_ (reference stress, 0.336 kPa). *τ*_tr_ is the native-stress filter time constant (0.15 s).

The published myofilament model provides active plus passive cell stress, *σ*_cell_, in kPa. Sigmoidal membrane-tension activation and context-dependent sensitivity have experimental support[68, 69]. Half activation occurs at a total sensed stress of 0.168 kPa. Whole-cell stress is a proxy for membrane tension; its mapping and the 0.15 s filter were not calibrated against tension measurements. Only native stress is filtered; the applied stimulus reaches the gate directly.

The availability, *h*, is modeled as:

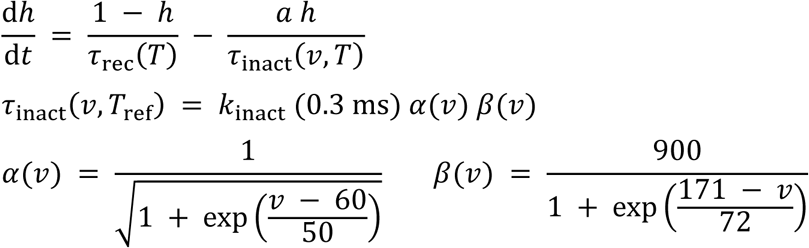

Here *v* is the numerical voltage in mV (*v* = 1000 *V_m_* with Vₘ in volts). T is absolute temperature (K), with *T*_ref_ = 295 K; *τ*_inact_ and *τ*_rec_ are inactivation and recovery time constants. *k*_inact_ is a dimensionless scale factor, and α and β are dimensionless voltage functions. Time constants are converted to seconds for integration. The voltage function follows the published model of Piezo1[43]. Recovery operates simultaneously, so its inactivation-term time scale differs from measured current decay. Extended description of the model is given in the **Supplemental Text1**. All model parameters for Piezo1 are listed in **Table S1**, the calibration conditions – in **Table S2** and **Fig. S5**, the test conditions – in **Table S3**.

## Author contributions

EE, JLH, MEP and WL designed the study. WL built the setup for combined optogenetic and ultrasound stimulation; WL performed the acoustic pressure simulations; MEP, CJC, YWH and OC conducted experiments related to optogenetic pacing and magnetic positioning; MEP and OC analyzed the data and prepared related figures; JLH optimized the Piezo1 delivery and characterization. JLH, YWH and WL conducted experiments related to combined optogenetic and ultrasound stimulation, analyzed data, and organized the related figures. MRP analyzed the Piezo1 expression in cardiac tissue and helped with figures. AF conducted computational simulations of hiPSC-CM responses to optogenetic and sonogenetic stimuli. MEP, JLH, WL and EE wrote the initial manuscript, with input from all authors.

## Acknowledgements

This work was supported in part by a grant from the National Institutes of Health R21EB026152 (EE) and from the Human Frontiers Science Program RGP010/2024 (EE). Biorender was used to create multiple figure panels.

## Data and materials availability

All experimental data are shown in the manuscript. The code for the computational modeling of optogenetic and sonogenetic responses in hiPSC-CMs will be deposited on Zenodo and released with the bioRxiv preprint. The reserved Zenodo DOI is https://doi.org/10.5281/zenodo.23430167; the code will also be released via GitHub (https://github.com/aminforouzandehmehr).

## REFERENCES

1. Choi, Y.S., et al., Fully implantable and bioresorbable cardiac pacemakers without leads or batteries. Nature Biotechnology, 2021. 39(10): p. 1228–1238.

2. Marchiano, S., et al., Gene editing to prevent ventricular arrhythmias associated with cardiomyocyte cell therapy. Cell Stem Cell, 2023. 30(5): p. 741.

3. Diagne, K., et al., Phase resetting in human stem cell derived cardiomyocytes explains complex cardiac arrhythmias. PLoS Comput Biol, 2026. 22(2): p. e1013935.

4. Kohut, A.R., et al., The potential of ultrasound in cardiac pacing and rhythm modulation. Expert Rev Med Devices, 2016. 13(9): p. 815–22.

5. Bilchick, K.C., et al., First-in-human noninvasive left ventricular ultrasound pacing: A potential screening tool for cardiac resynchronization therapy. Heart Rhythm O2, 2023. 4(2): p. 79–87.

6. Qu, J., et al., Expression and function of a biological pacemaker in canine heart. Circulation, 2003. 107(8): p. 1106–1109.

7. Hu, Y.-F., et al., Biological pacemaker created by minimally invasive somatic reprogramming in pigs with complete heart block. Science Translational Medicine, 2014. 6(245): p. 245ra94–245ra94.

8. Rosen, M.R., et al., Regenerative therapies in electrophysiology and pacing. J.Interv.Card Electrophysiol., 2008.

9. Plotnikov, A.N., et al., Biological pacemaker implanted in canine left bundle branch provides ventricular escape rhythms that have physiologically acceptable rates. Circulation, 2004. 109(4): p. 506–512.

10. Wang, J., et al., AAV-mediated long-term TBX18 expression causes cardiac fibrosis and fails to induce pacemaker activity in rodents. The Journal of Clinical Investigation, 2026. 136(13).

11. Zhang, T., et al., Human PSC-derived sinoatrial node-cardiac plexus assembloids model innervation-associated maturation of pacemaker systems. Cell Stem Cell, 2026. 33(6): p. 945–963.e11.

12. Emiliani, V., et al., Optogenetics for light control of biological systems. Nature Reviews Methods Primers, 2022. 2(1): p. 55.

13. Entcheva, E. and M.W. Kay, Cardiac optogenetics: a decade of enlightenment. Nat Rev Cardiol, 2021. 18(5): p. 349–367.

14. Ambrosi, C.M. and E. Entcheva, Optogenetics’ promise: pacing and cardioversion by light? Future Cardiol, 2014. 10(1): p. 1–4.

15. Hsueh, B., et al., Cardiogenic control of affective behavioural state. Nature, 2023. 615(7951): p. 292–299.

16. Alekseev, A., et al., Efficient and sustained optogenetic control of sensory and cardiac systems. Nat Biomed Eng, 2026. 10(2): p. 277–292.

17. Bruegmann, T., et al., Optogenetic control of heart muscle in vitro and in vivo. Nat Methods, 2010. 7(11): p. 897–900.

18. Arrenberg, A.B., et al., Optogenetic control of cardiac function. Science, 2010. 330(6006): p. 971–4.

19. Jia, Z., et al., Stimulating cardiac muscle by light: cardiac optogenetics by cell delivery. Circ Arrhythm Electrophysiol, 2011. 4(5): p. 753–60.

20. Riis, T. and J. Kubanek, Effective Ultrasonic Stimulation in Human Peripheral Nervous System. IEEE Trans Biomed Eng, 2022. 69(1): p. 15–22.

21. Kubanek, J., et al., Ultrasound Elicits Behavioral Responses through Mechanical Effects on Neurons and Ion Channels in a Simple Nervous System. J Neurosci, 2018. 38(12): p. 3081–3091.

22. Kubanek, J., et al., Ultrasound modulates ion channel currents. Sci Rep, 2016. 6: p. 24170.

23. Marquet, F., et al., Non-invasive cardiac pacing with image-guided focused ultrasound. Sci Rep, 2016. 6: p. 36534.

24. Gong, C., et al., A wearable non-invasive sonogenetic pacemaker. Nat Biomed Eng, 2026.

25. Huang, Y.S., et al., Sonogenetic Modulation of Cellular Activities in Mammalian Cells. Methods Mol Biol, 2021. 2312: p. 109–124.

26. Chua, C.J., et al., Integration of Engineered “Spark-Cell” Spheroids for Optical Pacing of Cardiac Tissue. Front Bioeng Biotechnol, 2021. 9: p. 658594.

27. Zhang, H., et al., Intramyocardial injection of allogeneic human induced pluripotent stem cell-derived cardiomyocytes in advanced ischemic heart failure: an early-stage randomized trial. Nature Medicine, 2026.

28. Hong, Y., et al., Pluripotent stem cell-derived cardiomyocyte transplantation: marching from bench to bedside. Sci China Life Sci, 2025.

29. Zhang, J.J., et al., Trials and tribulations of cell therapy for heart failure: an update on ongoing trials. Nat Rev Cardiol, 2025. 22(5): p. 372–385.

30. O’Hern, C., et al., Human heart-macrophage assembloids mimic immune-cardiac interactions and enable arrhythmia disease modeling. Cell Stem Cell, 2025. 32(11): p. 1671–1690.e13.

31. Song, Q., et al., Opto-SICM framework combines optogenetics with scanning ion conductance microscopy for probing cell-to-cell contacts. Commun Biol, 2023. 6(1): p. 1131.

32. Boyle, P.M., et al., OptoGap is an optogenetics-enabled assay for quantification of cell-cell coupling in multicellular cardiac tissue. Sci Rep, 2021. 11(1): p. 9310.

33. Kostecki, G.M., et al., Optogenetic current in myofibroblasts acutely alters electrophysiology and conduction of co-cultured cardiomyocytes. Sci Rep, 2021. 11(1): p. 4430.

34. Yu, J. and E. Entcheva, Inscribing Optical Excitability to Non-Excitable Cardiac Cells: Viral Delivery of Optogenetic Tools in Primary Cardiac Fibroblasts. Methods Mol Biol, 2016. 1408: p. 303–17.

35. Syeda, R., et al., Piezo1 Channels Are Inherently Mechanosensitive. Cell Rep, 2016. 17(7): p. 1739–1746.

36. Zhu, J., et al., The mechanosensitive ion channel Piezo1 contributes to ultrasound neuromodulation. Proc Natl Acad Sci U S A, 2023. 120(18): p. e2300291120.

37. Klimas, A., et al., Multimodal on-axis platform for all-optical electrophysiology with near-infrared probes in human stem-cell-derived cardiomyocytes. Prog Biophys Mol Biol, 2020. 154: p. 62–70.

38. Klimas, A., et al., OptoDyCE as an automated system for high-throughput all-optical dynamic cardiac electrophysiology. Nat Commun, 2016. 7: p. 11542.

39. Datta, P., et al., Recent Advances and Future Directions in Sonodynamic Therapy for Cancer Treatment. BME Frontiers, 2024. 5: p. 0080.

40. Qiu, Z., et al., The Mechanosensitive Ion Channel Piezo1 Significantly Mediates In Vitro Ultrasonic Stimulation of Neurons. iScience, 2019. 21: p. 448–457.

41. Li, W., et al., Response of human iPSC-cardiomyocytes to adrenergic drugs assessed by high-throughput pericellular oxygen measurements and computational modeling. J Mol Cell Cardiol Plus, 2026. 15: p. 100834.

42. Williams, J.C., et al., Computational optogenetics: empirically-derived voltage- and light-sensitive channelrhodopsin-2 model. PLoS Comput Biol, 2013. 9(9): p. e1003220.

43. Forouzandehmehr, M., et al., Mechanosensitive Channel Piezo1 in R403Q Hypertrophic Cardiomyopathy: A Computational Study. 2023 Computing in Cardiology (CinC), 2023. 50: p. 1–4.

44. Williams, J.C. and E. Entcheva, Optogenetic versus Electrical Stimulation of Human Cardiomyocytes: Modeling Insights. Biophys J, 2015. 108(8): p. 1934–45.

45. Coste, B., et al., Piezo1 ion channel pore properties are dictated by C-terminal region. Nat Commun, 2015. 6: p. 7223.

46. Heinson, Y.W., J.L. Han, and E. Entcheva, OptoDyCE-plate as an affordable high throughput imager for all optical cardiac electrophysiology. Journal of Molecular and Cellular Cardiology Plus, 2023. 6: p. 100054.

47. Voges, H.K., et al., Vascular cells improve functionality of human cardiac organoids. Cell Reports, 2023. 42(5).

48. Wu, X., et al., Reimagining human-centric drug development with new approach methodologies. Science, 2026. 392(6796): p. 371–378.

49. Jebran, A.F., et al., Engineered heart muscle allografts for heart repair in primates and humans. Nature, 2025. 639(8054): p. 503–511.

50. Ivanovski, F., et al., Ultrasound-mediated spatial and temporal control of engineered cells in vivo. Nature Communications, 2024. 15(1): p. 7369.

51. Hu, H., et al., A wearable cardiac ultrasound imager. Nature, 2023. 613(7945): p. 667–675.

52. Sugiura, T., D.C. Shahannaz, and B.E. Ferrell, Current Status of Cardiac Regenerative Therapy Using Induced Pluripotent Stem Cells. Int J Mol Sci, 2024. 25(11).

53. Shen, X., et al., Sensitization of nerve cells to ultrasound stimulation through Piezo1-targeted microbubbles. Ultrason Sonochem, 2021. 73: p. 105494.

54. Coste, B., et al., Piezo1 and Piezo2 are essential components of distinct mechanically activated cation channels. Science, 2010. 330(6000): p. 55–60.

55. Duque, M., et al., Sonogenetic control of mammalian cells using exogenous Transient Receptor Potential A1 channels. Nat Commun, 2022. 13(1): p. 600.

56. Ambrosi, C.M., et al., Adeno-Associated Virus Mediated Gene Delivery: Implications for Scalable in vitro and in vivo Cardiac Optogenetic Models. Front Physiol, 2019. 10: p. 168.

57. Siquara da Rocha, L.O., et al., Mapping Cell-in-Cell Structures in Oral Squamous Cell Carcinoma. Cells, 2023. 12(19).

58. Baillargeon, P., et al., Automating a Magnetic 3D Spheroid Model Technology for High-Throughput Screening. SLAS Technol, 2019. 24(4): p. 420–428.

59. Pozo, M.R., et al., Control of cardiac waves in human iPSC-CM syncytia by a Halbach array and magnetic nanoparticles. Biophys J, 2025. 124(8): p. 1273– 1284.

60. Li, W., J.L. Han, and E. Entcheva, Protein and mRNA Quantification in Small Samples of Human-Induced Pluripotent Stem Cell-Derived Cardiomyocytes in 96-Well Microplates. Methods Mol Biol, 2022. 2485: p. 15–37.

61. GTExConsortium, The GTEx Consortium atlas of genetic regulatory effects across human tissues. Science, 2020. 369(6509): p. 1318–1330.

62. Pozo, M.R., et al., Comparative transcriptomics analysis of histone deacetylases, transcription factors, and ion channel genes in human iPSC-cardiomyocytes vs. the adult human heart. The Journal of Precision Medicine: Health and Disease, 2026. 6: p. 100031.

63. Pressler, M.P., A. Horvath, and E. Entcheva, Sex-dependent transcription of cardiac electrophysiology and links to acetylation modifiers based on the GTEx database. Front Cardiovasc Med, 2022. 9: p. 941890.

64. Andrew, H., Voltage to pressure conversion: are you getting ‘phased’ by the problem? Journal of Physics: Conference Series, 2004. 1(1): p. 57.

65. Wu, J., et al., Inactivation of Mechanically Activated Piezo1 Ion Channels Is Determined by the C-Terminal Extracellular Domain and the Inner Pore Helix. Cell Rep, 2017. 21(9): p. 2357–2366.

66. Zheng, W., E.O. Gracheva, and S.N. Bagriantsev, A hydrophobic gate in the inner pore helix is the major determinant of inactivation in mechanosensitive Piezo channels. Elife, 2019. 8.

67. Lewis, A.H., et al., Transduction of Repetitive Mechanical Stimuli by Piezo1 and Piezo2 Ion Channels. Cell Rep, 2017. 19(12): p. 2572–2585.

68. Lewis, A.H. and J. Grandl, Mechanical sensitivity of Piezo1 ion channels can be tuned by cellular membrane tension. Elife, 2015. 4.

69. Cox, C.D., et al., Removal of the mechanoprotective influence of the cytoskeleton reveals PIEZO1 is gated by bilayer tension. Nat Commun, 2016. 7: p. 10366.

