## Supplement for "Optogenetic and sonogenetic pacing of unmodified cardiac tissue upon quick integration with specialized biological actuators"

### SUPPLEMENTARY INFORMATION
### for

#### Optogenetic and sonogenetic pacing of unmodified cardiac tissue upon quick integration with specialized biological actuators

**Morgan E. Pettebone[1*], Julie L. Han[1*],**
Amin Forouzandehmehr[2], Olivia Chesla[1], Christianne J. Chua[1], Maria R. Pozo[1], Yuli W. Heinson[1], Wei Liu[1,3], Emilia Entcheva[1#]

[1]Department of Biomedical Engineering, School of Engineering and Applied Science, The George Washington University, Washington DC, USA

[2]Faculty of Medicine and Health Technology, Tampere University, Finland
[3]Optical Imaging Laboratory, Harbin Institute of Technology, Shenzhen, China

***Short title:*** *Controlling cardiac tissue with quickly integrating biological actuators*

*These authors contributed equally to the work

**# Corresponding author:**

Emilia Entcheva, PhD

Department of Biomedical Engineering
George Washington University
800 22nd St NW, Suite 5000
Washington, DC 20052

**Figure S1.** iPSC-CM monolayer entrainment ranges after 2 hrs of integration using ChR2-HEK spheroids mixed with several types of mNPs (n=5 monolayers per mNP type). All ChR2-HEK spheroids are 10k cells in size.

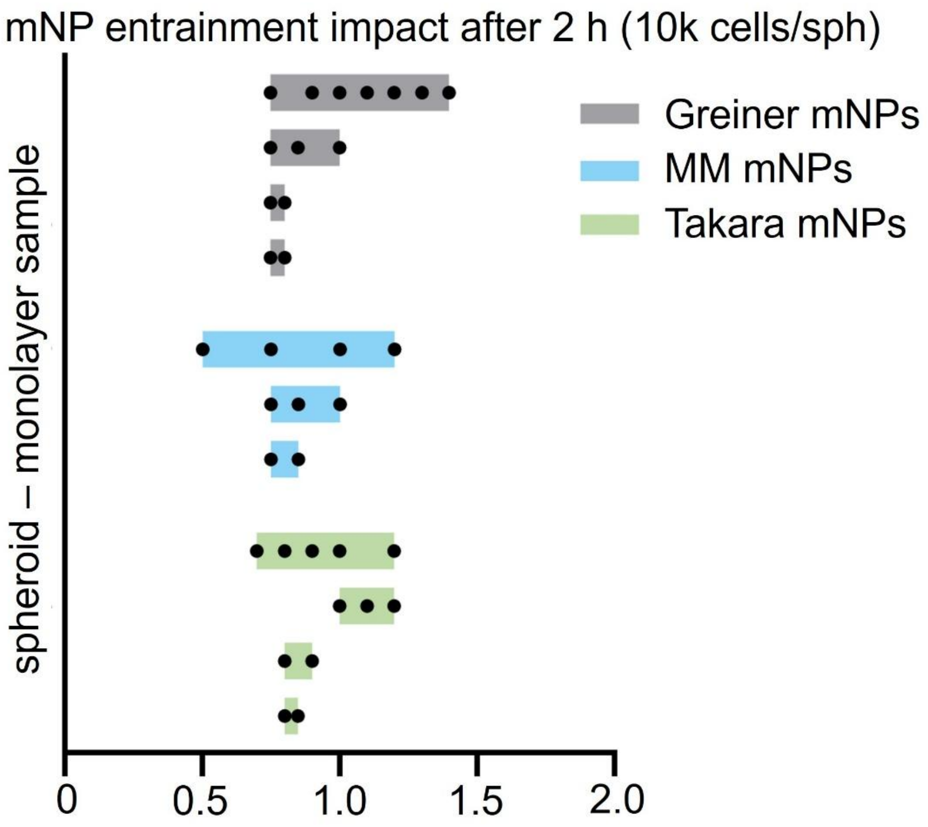

**Figure S2** iPSC-CM spheroid entrainment ranges after 1 hr integration with ChR2-HEK spheroids (n= 19 spheroid pairs probed). Both spheroids are 1k cells in size.

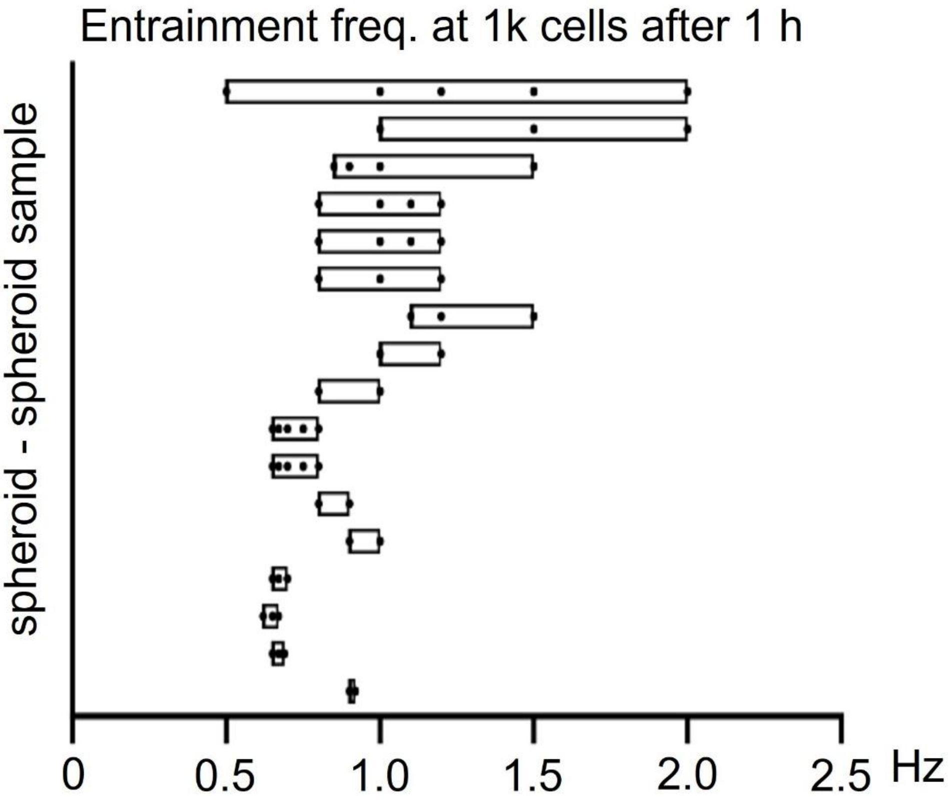

**Figure S3.** iPSC-CM spheroid entrainment ranges after 2 hr integration with ChR2-HEK spheroids (n= 8 spheroid pairs probed). Both spheroids are 5k cells in size.

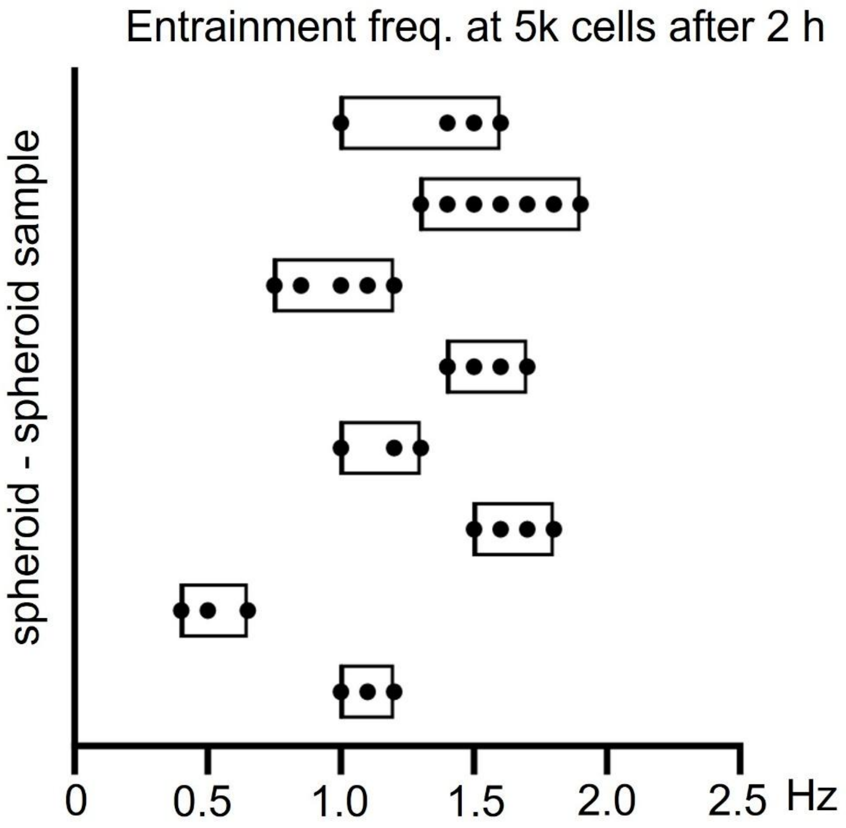

**Figure S4. A.** Quantification of relative Piezo1 levels in iPSC-CMs after combinations of MOI and viral dosing order of Ad-ChR2 and Ad-Piezo1. **B.** Quantification of normalized Piezo1 in human cFBs after combinations of MOI and viral dosing order of Ad-Piezo1 and Ad-ChR2. Sequential treatments were 24h apart.

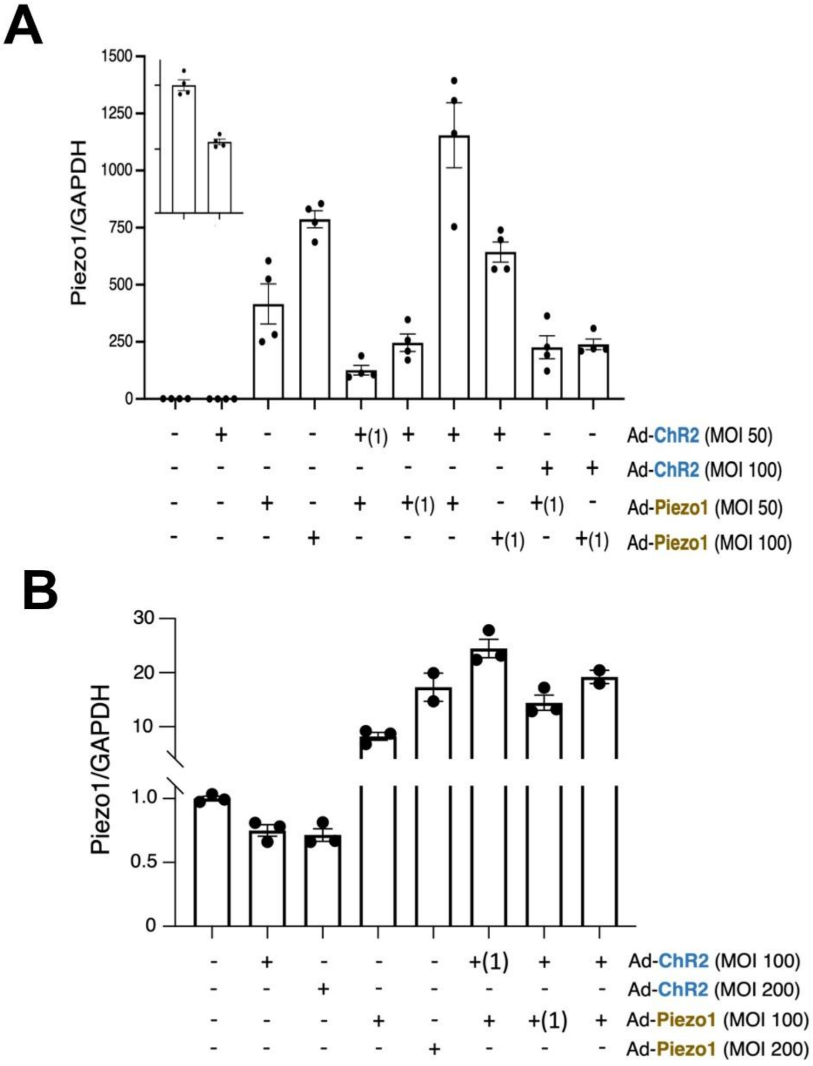

**Figure S5.** Fitting the decay time constant against voltage for Piezo 1 at the nominal recording temperature of 295 K. Experimental data used to fit the model curve (in teal) are indicated in the legend; also data points not used for the fit are overlaid (see "check" in the legend). Error bars for fitting targets are approximate values digitized from the source figure; the Albuisson et al. 2013 [7] human Piezo1 data show mean±2 SEM. The Wu et al. 2017 [4] additional (check) data are reported as mean values without error bars.

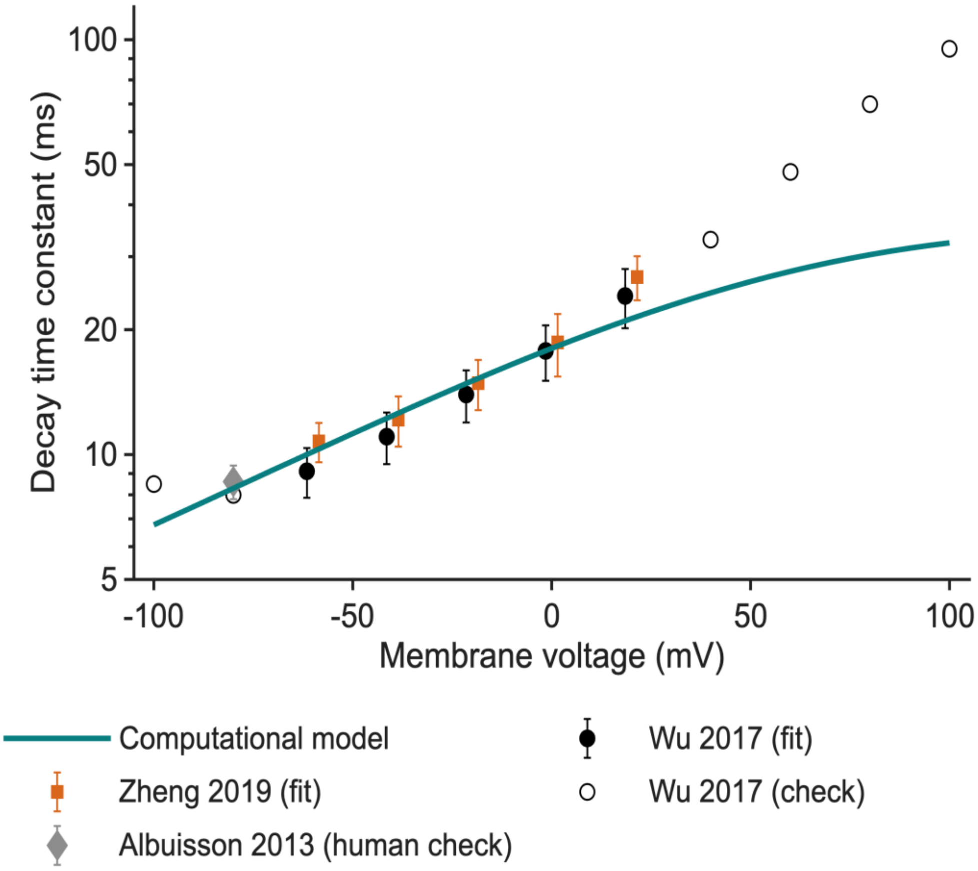

**Suppl. Table S1. Computational model parameters for Piezo1 used to generate Fig. 8A**

| Parameter | Value | Basis |
|---|---|---|
| $g_P$ | 91.17 S/F | Effective channel number; 300 × released conductance [3]; nominal unitary conductance [8] |
| $\sigma_{\mathrm{ref}}$ | 0.336 kPa | Application setting; released reference 0.056 kPa [3]; new value not measured |
| $\tau_{\mathrm{tr}}$ | 0.150 s | Assumed native-stress transmission; no quantitative experimental calibration |
| $\tau_a$ | 3.162 ms | Fitted to fast deactivation [6]; equal activation time assumed |
| $k_{\mathrm{inact}}$ | 0.9098 | Fitted to voltage-dependent decay [4,5] |
| $\tau_{\mathrm{rec}}$ | 0.2023 s | Fitted to approximate half recovery [6] |
| $f_{\mathrm{Ca}}$ | 0.05 | Approximate constant-field Na/Ca current share [8,9]; fixed allocation assumed |
| $E_P$ | 0 mV | Inherited near-zero reversal [3]; bidirectional permeation is experimentally supported [9,10] |

Kinetic time constants are specified at $T_{\mathrm{ref}}$ = 295 K and scaled to the simulation temperature of 304 K. The 0.15 s mechanical filter is not temperature-scaled.

**Suppl. Table S2. Calibration of the computational model of Piezo1**

| Calibration target | Source and preparation | Targets | Assessment |
|---|---|---|---|
| Decay at −60, −40, −20, 0 and +20 mV | Wu 2017 [4]; mouse, whole cell | 5 | 5/5 within digitized error bars |
| Decay at the same five voltages | Zheng 2019 [5]; mouse, whole cell | 5 | 4/5 within digitized error bars |
| Fast deactivation after a 40 ms step | Lewis 2017 [6]; mouse, cell attached, −80 mV | 1 | 4.0 ms target; 3.86 ms model; assigned ±20% tolerance |
| Half recovery after paired 100 ms steps | Lewis 2017 [6]; mouse, cell attached, −80 mV | 1 | 0.140 s target; 0.144 s model; assigned ±20% tolerance |

**Suppl. Table S3. Model testing outside of the calibration parameters for Piezo1**

| Check | Experimental target | Model | Source |
|---|---|---|---|
| Human decay at −80 mV | 8.6 ± 0.4 ms (SEM) | 8.33 ms | Albuisson 2013 [7] |
| Residual after 300 ms at −80 mV | 0.096 ± 0.040 (SD) | 0.0598 | Lewis 2017 [6] |
| Mouse decay at −80 mV | 11.9 ± 0.6 ms (SEM) | 8.29 ms | Zheng 2019 [5] |
| Mouse decay at −100 / −80 mV | Approximately 8.5 / 8.0 ms | 6.77 / 8.29 ms | Wu 2017 [4] |
| Mouse decay at +40 to +100 mV | Approximately 33–95 ms | 24.5–32.4 ms | Wu 2017 [4] |

**Supplementary Text1:**

#### Computational modeling of optogenetic and sonogenetic stimulation of human iPSC-CMs with expression of ChR2 and Piezo1

For the simulations we used our published model of human iPSC-CMs - hiMCES2026 [1], with inserted ChR2 model of Williams et al. [2] and a modified Piezo1 model based on our published CinC2023 formulation [3]. Simulations were performed at 31 °C; experiments were conducted at room temperature. Intrinsic host parameters were retained; ChR2 and Piezo1 currents were added to the model. For the Piezo1 description, the voltage-sensing gate from [3] was replaced by an availability gate, and three kinetic parameters were fitted to published channel recordings [4–6]. Effective functional Piezo1 channel number per cell was represented by a whole-cell conductance of 9 nS, corresponding to approximately 310 channels using a nominal unitary conductance of 29.1 pS [8]. This functional channel number permitted mechanical capture. It should be noted that this is a conductance-equivalent estimate, not a measurement of protein expression or surface density. Native cellular tension reached the gate through a first-order filter with a 0.15 s time constant; the imposed mechanical stimulus acted directly on the gate.

In the simulations, cells were conditioned by 0.5 Hz optogenetic pacing (10 ms at 0.15 mW/mm²). The responses to three optical pulses are shown, followed 1 s later by one 0.5 s mechanical stress pulse of 0.336 kPa representing the US stimulus. No acoustic propagation or conversion from acoustic intensity to gate stress was modeled. Equations were integrated with ode15s in MATLAB R2026a.

##### Piezo1 current and mechanical activation

$$I_{\text{Piezo1}} = g_P\, a\, h\, (V_m - E_P), \quad E_P = 0\,\text{V}$$

Current is normalized to membrane capacitance (A/F), conductance is in S/F, and voltage in this current equation is in volts. The activation gate a and availability h are dimensionless.

$$\frac{\mathrm{d}a}{\mathrm{d}t} = \frac{a_\infty - a}{\tau_a(T)}$$

$$a_\infty = \frac{1}{1 + \exp\left(-\dfrac{\dfrac{5\sigma}{\sigma_{\text{ref}}} - 2.5}{0.25}\right)}$$

$$\sigma = \sigma_{\text{cell,s}} + \sigma_{\text{stim}}$$

$$\frac{\mathrm{d}\sigma_{\text{cell,s}}}{\mathrm{d}t} = \frac{\sigma_{\text{cell}} - \sigma_{\text{cell,s}}}{\tau_{\text{tr}}}$$

The published myofilament model provides active plus passive cell stress, $\sigma_{\text{cell}}$, in kPa. Sigmoidal membrane-tension activation and context-dependent sensitivity have experimental support [12,13]. Whole-cell stress is a proxy for membrane tension; its mapping and the 0.15 s filter were not calibrated against tension measurements. Only native stress is filtered; the applied stimulus reaches the gate directly. Half activation occurs at a total sensed stress of 0.168 kPa.

##### Channel kinetics and parameters

###### Availability gate

$$\frac{\mathrm{d}h}{\mathrm{d}t} = \frac{1 - h}{\tau_{\text{rec}}(T)} - \frac{a\,h}{\tau_{\text{inact}}(v, T)}$$

$$\tau_{\text{inact}}(v, T_{\text{ref}}) = k_{\text{inact}}\,(0.3\text{ ms})\,\alpha(v)\,\beta(v)$$

$$\alpha(v) = \frac{1}{\sqrt{1 + \exp\left(\frac{v - 60}{50}\right)}} \qquad \beta(v) = \frac{900}{1 + \exp\left(\frac{171 - v}{72}\right)}$$

Here v is the numerical voltage in mV ($v = 1000\,V_m$ with $V_m$ in volts). The voltage function follows the published model of Piezo1 [3]. Recovery operates simultaneously, so its inactivation-term time scale differs from measured current decay. All model parameters for Piezo1 are listed in **Table S1**.

###### Temperature dependence

$$\tau_j(T) = \tau_j(T_{\text{ref}})\,2^{\frac{T_{\text{ref}} - T}{10\text{ K}}}$$

This scaling applies to activation, inactivation and recovery, using a nominal recording reference of 295 K (approximately 22 °C). Cooling-induced slowing of Piezo1 inactivation is experimentally supported [11], but a common $Q_{10}$ of 2 for these three processes is an assumption, not a measured value. It gives a factor of 0.5359 at 304 K. The mechanical transmission time is not temperature-scaled.

##### Functional channel number and ionic currents

###### Meaning of channel number

The conductance-normalized representation can be expressed as an effective number of functional channels. With the published host capacitance of 98.71 pF [1] and the selected conductance,

$$N_{\text{eff}} = \frac{g_P\,C_m}{\gamma} = \frac{9.00\text{ nS}}{29.1\text{ pS}} \approx 310.$$

Coste et al. measured a wild-type mouse Piezo1 unitary conductance of 29.1 ± 0.4 pS (SEM) under their recording conditions [8]. This is used only as a nominal conversion: unitary conductance depends on solutions and temperature [8,9,11]. The number refers to the available functional population, not the number open at rest; its instantaneous open fraction is (a·h). Channel density would require a stated normalization to capacitance or membrane area.

###### Simplified ion accounting

Net inward Piezo1 current is allocated 95% to $Na^+$ and 5% to $Ca^{2+}$. The approximate $Ca^{2+}$ fraction is motivated by Goldman–Hodgkin–Katz (GHK) constant-field currents near −75 mV. Coste et al. reported $\frac{P_{\text{Ca}}}{P_{\text{Cs}}} = 1.21$ for mouse Piezo1 [8].

Gnanasambandam et al. reported human $P_K : P_{\text{Cs}} : P_{\text{Na}} : P_{\text{Li}} = 1 : 0.88 : 0.82 : 0.71$ [9], giving $\frac{P_{\text{Cs}}}{P_{\text{Na}}} \approx 1.07$. Combining these cross-species values gives:

$$\frac{P_{\text{Ca}}}{P_{\text{Na}}} = \frac{P_{\text{Ca}}}{P_{\text{Cs}}} \frac{P_{\text{Cs}}}{P_{\text{Na}}} \approx 1.21 \times 1.07 \approx 1.30.$$

For an illustrative constant-field calculation at −75 mV and 304 K, using the host Tyrode extracellular $Na^+$ and $Ca^{2+}$ concentrations (135 and 1.33 mM [1]), intracellular $Na^+$ of 10 mM and $Ca^{2+}$ of 0.1 μM, the $Ca^{2+}$ share of the combined inward $Na^+$ and $Ca^{2+}$ currents is approximately 4.6%, rounded to 5%:

$$f_{\text{Ca,GHK}} = \frac{|I_{\text{Ca}}|}{|I_{\text{Na}}| + |I_{\text{Ca}}|} \approx 0.046.$$

The 5% value is therefore a derived approximation, not a measured fraction of total Piezo1 current. In this model it is kept fixed and applied to net inward current; $K^+$ counterflow and voltage-dependent changes in ionic fractions are not resolved. Net outward current is assigned to $K^+$ efflux with fixed intracellular $K^+$. Simultaneous inward $Na^+$/$Ca^{2+}$ flux during net outward current is omitted. ChR2 enters voltage but no ion balance. Calcium results are qualitative. Piezo1-mediated $Ca^{2+}$ entry and stretch-evoked $Ca^{2+}$ signaling in mouse cardiomyocytes [14] support the qualitative pathway.

**Calibration and checks**

The sources and conditions for model calibration for Piezo1 are listed in **Table S2,** see also **Figure S5**. **Table S3** lists conditions outside the calibration data that were used for testing. The three kinetic parameters were fitted to 12 targets from heterologously expressed mouse Piezo1, combining whole-cell probe stimulation and cell-attached pressure-clamp recordings. The ten voltage-dependent decay targets and their error bars were digitized from the CinC2023 figure [3], which reproduced Wu et al. [4] and Zheng et al. [5]. These are approximate digitized targets rather than raw recordings. The deactivation target is the fast component of a two-phase tail; the recovery target is an approximate half-recovery delay interpolated from the shallow portion of the recovery curve [6]. The isolated-channel evaluator used a 1 ms stimulus rise for whole-cell decay and a 2 ms rise/withdrawal for pressure-clamp protocols, with saturating effective stress. Decay was measured by fitting an exponential plus offset from the peak. Normalized conductance (a·h) was used to isolate gating kinetics, including at 0 mV where the ohmic current is zero. Deactivation was fitted for 30 ms after release; paired-pulse recovery used baseline-subtracted peak ratios. These waveform approximations are modeling assumptions.

Nine of ten fitted decay points lie within their digitized error bars. The exception is Zheng at +20 mV: 26.8 ± 3.2 ms versus 21.3 ms in the model. The two other fitted targets meet their assigned 20% tolerances, which are not experimental error bars.

**In Figure S5,** we overlaid data from human Piezo1 (150 ms stimulus was used there) after the fitting. The high-voltage Wu values were also overlaid post-fitting. The full recovery curve is partly calibrated because its half-recovery delay entered the fit. Recovery and train limitations are stated below.

#### Known model limitations

This version of the model misses slow recovery phases and underestimates current depression during trains of brief stimuli, particularly at 0.5–5 Hz [6]. It also misses several unfitted mouse decay values (**Table S3**). Its single availability gate is a phenomenological approximation. The native-stress filter, conductance scale and stress normalization represent assumptions rather than measurements.

The assumed Piezo1 reversal potential is 0 mV. During the mechanical pulse, positive membrane voltage therefore produces an outward current, reaching +0.41 A/F just after the upstroke. Outward PIEZO1 currents at positive voltages have been recorded experimentally [9,10], i.e. these are biophysically plausible. The amplitude and time course in **Figure 8A** have not been validated directly against a cardiomyocyte recording. This single-cell simulation does not model acoustic propagation, actuator spheroids or coupling to unmodified cardiac tissue, and it does not establish an acoustic stimulation threshold.

#### Simulation record

Both the model and the Piezo1-off control met the selected conditioning criteria after 1000 s at 304 K. The model had APD90 618 ms and peak $Ca^{2+}$ 189 nM; control APD90 was 616 ms. Between 200 s blocks, relative changes in APD90, peak $Ca^{2+}$ and intracellular $Na^{+}$ were below $2 \times 10^{-4}$, and absolute gate changes were below $10^{-3}$.

All three 10 ms optical pulses captured at 0.15 mW/mm². The lowest tested capturing mechanical amplitude was 0.168 kPa; 0.119 kPa did not capture. Capture therefore occurred at a threshold bracketed between these sampled amplitudes. Every larger tested amplitude through 1.344 kPa captured. Figure 8A uses 0.336 kPa, approximately 3.4 times the peak native cell stress of 0.0978 kPa. The no-stimulus control and the Piezo1-off control had no upstroke in the mechanical-stimulus assessment window (5.0–5.6 s).

The optical overlay uses the second light beat (APD90 624 ms); mechanical APD90 is 582 ms. Peak currents are −2.19 A/F for ChR2 and −3.68 A/F inward / +0.41 A/F outward for Piezo1. Mechanical peak $Ca^{2+}$ is 147 nM versus 189 nM during light pacing; the first no-stimulus spontaneous beat reaches 143 nM.

#### References


[1] Li W, et al. Response of human iPSC-cardiomyocytes to adrenergic drugs assessed by high-throughput pericellular oxygen measurements and computational modeling. Journal of Molecular and Cellular Cardiology Plus. 2026;15:100834.

[2] Williams JC, et al. Computational Optogenetics: Empirically-Derived Voltage- and Light-Sensitive Channelrhodopsin-2 Model. PLoS Computational Biology. 2013;9:e1003220.

[3] Forouzandehmehr M, et al. Mechanosensitive Channel Piezo1 in R403Q Hypertrophic Cardiomyopathy: A Computational Study. Computing in Cardiology. 2023;50. https://doi.org/10.22489/CinC.2023.359.

[4] Wu J, et al. Inactivation of Mechanically Activated Piezo1 Ion Channels Is Determined by the C-Terminal Extracellular Domain and the Inner Pore Helix. Cell Reports. 2017;21:2357–2366.

[5] Zheng W, Gracheva EO, Bagriantsev SN. A hydrophobic gate in the inner pore helix is the major determinant of inactivation in mechanosensitive Piezo channels. eLife. 2019;8:e44003.

[6] Lewis AH, et al. Transduction of repetitive mechanical stimuli by Piezo1 and Piezo2 ion channels. Cell Reports. 2017;19:2572–2585.

[7] Albuisson J, et al. Dehydrated hereditary stomatocytosis linked to gain-of-function mutations in mechanically activated PIEZO1 ion channels. Nature Communications. 2013;4:1884.

[8] Coste B, et al. Piezo1 ion channel pore properties are dictated by C-terminal region. Nature Communications. 2015;6:7223.

[9] Gnanasambandam R, Bae C, Gottlieb PA, Sachs F. Ionic Selectivity and Permeation Properties of Human PIEZO1 Channels. PLoS One. 2015;10:e0125503.

[10] Moroni M, Servin-Vences MR, Fleischer R, Sánchez-Carranza O, Lewin GR. Voltage gating of mechanosensitive PIEZO channels. Nature Communications. 2018;9:1096.

[11] Zheng W, Nikolaev YA, Gracheva EO, Bagriantsev SN. Piezo2 integrates mechanical and thermal cues in vertebrate mechanoreceptors. Proceedings of the National Academy of Sciences USA. 2019;116:17547–17555.

[12] Lewis AH, Grandl J. Mechanical sensitivity of Piezo1 ion channels can be tuned by cellular membrane tension. eLife. 2015;4:e12088.

[13] Cox CD, et al. Removal of the mechanoprotective influence of the cytoskeleton reveals PIEZO1 is gated by bilayer tension. Nature Communications. 2016;7:10366.

[14] Jiang F, et al. The mechanosensitive Piezo1 channel mediates heart mechano-chemo transduction. Nature Communications. 2021;12:869.
